# A lobule-specific neuronal representation of song temporal structure in the songbird cerebellum

**DOI:** 10.1101/2025.11.10.687662

**Authors:** Roman Ursu, Eduarda Centeno, Arthur Leblois

## Abstract

The cerebellum is involved in the acquisition and production of speech as revealed by clinical evidence and imaging studies, but its specific role however remains unclear. Songbirds provide a unique model to study the neural mechanisms of speech learning and production. Recent evidence highlights a contribution of the cerebellum to syllable duration in songbirds. Here, we aim at better understanding how and which cerebellar circuits may contribute to the tight control of syllable duration in zebra finches. We first confirmed that lesions in the lateral cerebellum affect syllable duration. We then recorded neuronal activity in the various cerebellar lobules in response to song playback and during singing with or without auditory feedback perturbation. We found that many cerebellar lobules receive non-selective auditory information locked to syllable boundaries during song playback, both in anaesthetized and awake behaving birds. During singing, cerebellar neurons in several lobules display song-locked responses with varying degrees of alignment to their playback responses and no alteration during auditory feedback perturbation. Singing-related activity tends to correlate with the fluctuations in syllable duration rather than any acoustic song feature. Importantly, neurons in lobule IV of the cerebellum are largely unaffected by auditory stimulation but display syllable-locked firing rate modulations during singing that precisely encode syllable boundaries with a sharp and tight increase in firing at syllable onsets and offsets. Altogether, our findings reveal cerebellar signals that may contribute to the control of the duration of vocal elements during singing, possibly serving as a forward model of song temporal features.

## Introduction

Speech acquisition and production are complex sensorimotor behaviors that rely on multiple brain structures, including the cerebral cortex, thalamus, basal ganglia, and cerebellum (Lieberman, 2002; Vargha-Khadem et al., 2005; Ziegler & Ackermann, 2017). Cerebellar dysfunction disrupts speech production (Ackermann & Ziegler, 1991; Darley et al., 1969; Fiez, 2016), perception (Ackermann et al., 1992, 1997; Gandour et al., 1988) and learning (Desmond & Fiez, 1998; Leiner et al., 1991; Muehlbacher et al., 2025). Imagine studies localize speech-related cerebellar activity to lateral hemispheric regions (Ackermann & Brendel, 2016). Specifically, lobules V and VI as well as Crus I/II appear to contribute to motor control of speech (Callan et al., 2007; Frings et al., 2006; Stoodley et al., 2012; Thürling et al., 2011) whereas lobules VII, VIII, and more recently IX have been implicated in language comprehension and verbal memory (Jobson et al., 2024; Tourville et al., 2008). Nonetheless, the precise functional roles of these cerebellar subregions in speech and the associated neural mechanisms remain poorly understood (Mariën et al., 2014; Silveri, 2021).

Songbirds provide a powerful model for studying the neural bases of speech learning and production (Brainard & Doupe, 2013). Juveniles learn their song through a process analogous to human speech learning, suggesting that similar neural mechanisms may be involved (Brainard & Doupe, 2002; Doupe & Kuhl, 1999). Songbirds feature a dedicated neuronal song circuit to produce and learn the song (Mooney, 2009). This song circuit includes a basal ganglia–thalamo–cortical loop homologous to circuits supporting human speech (Doupe et al., 2005; Jarvis, 2019). Recent evidence revealed a functional pathway from the cerebellum to both the song-related basal ganglia and the song production cortical circuit (Nicholson et al., 2018; Pidoux et al., 2018), similar to the cerebello-thalamo-basal ganglia and cerebello-thalamo-cortical pathways in primates (Bostan et al., 2010). Lesions of the deep cerebellar nuclei (DCN) in adults produce lasting temporal disruptions in song (Radic et al., 2024). In juveniles, lesions in the lateral cerebellar nucleus interferes with song learning, particularly the maturation of temporal features (Pidoux et al., 2018). Despite this evidence, the cerebellum’s exact role in song production and learning, and the underlying neural computations, remain unresolved.

Prevailing theories of cerebellar function in vertebrates often invoke internal models (Ebner & Pasalar, 2008; Wolpert & Kawato, 1998). In this framework, the cerebellum integrates an efference copy of motor commands with concurrent sensory feedback to compute discrepancies between predicted and actual inputs (Cullen, 2023; Lisberger et al., 1987). These discrepancies drive real-time corrections and update internal models, with particular importance for timing. Temporal precision is a key domain for these corrections (Bareš et al., 2019; Ivry et al., 1988; Okada et al., 2022), as adjustments often occur through fine-tuning the timing of muscle activation (Becker & Person, 2019; Vilis & Hore, 1980).

Where and how motor and auditory signals related to song and its temporal features are represented in the avian cerebellum remains unknown. While neurons in lobules II, V, VI, VII, and VIII respond to auditory input in anesthetized birds (Gross, 1970; Whitlock, 1952), the nature of auditory signals reaching the cerebellum of awake, singing birds is less clear. To investigate this, we recorded neuronal activity in the lateral cerebellum of zebra finches during spontaneous singing (with and without distorted auditory feedback) and during song playback. Our results reveal (i) widespread, nonselective auditory responses across the cerebellum in both anesthetized and awake birds, and (ii) strong singing-related modulation in multiple lobules. This modulation sometimes overlapped with playback-evoked responses but was largely unaffected by auditory feedback perturbations. These findings highlight cerebellar signals that may contribute to the implementation of internal models during vocal production.

## Results

### Lesion of the lateral DCN reduce syllable and motif duration

Previous evidence revealed that DCN lesions can affect syllable duration both during song learning (Pidoux et al., 2018) and in adult zebra finches (Radic et al., 2024). As the latter study reports effects of lesions which spread across all DCNs, we first confirmed that lesion restricted to the lateral DCN (latDCN) systematically affects syllable duration. Electrolytic (n=2/9 birds) or chemical (Ibotenic acid, n=7/9 birds) lesions were performed in the latDCN bilaterally (see histological control in Suppl Fig 1A). As shown in Fig. 1A-C for an example syllable, latDCN lesion reduced syllable duration (related t-test, p<0.001, all renditions across multiple days pre and post-lesion on Fig. 1C). Over all lesioned birds, syllable duration was significantly decreased (Fig. 1D, 141 +/- 80 ms before lesion vs 135 +/-79 ms after lesion, n=33 motif syllables from 9 birds, related t-test on log data, p< 0.001, see Methods). Gaps between syllables displayed inconsistent changes in duration within and across birds, overall showing no significant difference (Fig. 1E, 35 +/- 14 ms vs 36 +/- 14 ms, n=23 gaps in 9 birds, related t-test, p>0.05). As a consequence, song motif duration significantly decreased after lesion (567 +/- 187 ms vs 553 +/-183 ms, n=9 motifs from 9 birds, related t-test, p<0.05, Fig. 1F). On the contrary, sham lesions did not induce any change in syllable duration (116 +/- 72 ms vs 116 +/- 72 ms, p>0.05, see Suppl Fig 1B), and neither in gap (34 +/- 17 ms vs 33 +/- 17 ms, p>0.05) or motif duration (633 +/- 94 ms vs 622 +/- 91 ms, p>0.05). Altogether, syllable duration strongly correlates with lesion size across birds (Spearman test, p<0.05, Suppl Fig 1C). LatDCN lesions also produced a decrease in syllable amplitude (Fig. 1G, 0.37 +/- 0.65 a.u. vs 0.32 +/- 0.59 a.u., related t-test on log data, p<0.001, see Methods). As changes in amplitude can affect syllable duration measure, we confirmed that the decrease in syllable duration is still significant after compensation for changes in amplitude (Suppl Fig 1D-E, 141 +/- 80 ms vs 138 +/- 78 ms, related t-test on log data, p< 0.001, see Methods). Overall, our results confirm that latDCN lesions strongly affect song temporal features in zebra finches, with a significant decrease in syllable and motif duration. Thus, the lateral cerebellum participates in the tight control of syllable timing, not solely during song learning but also during singing in adulthood.

**Figure 1:**
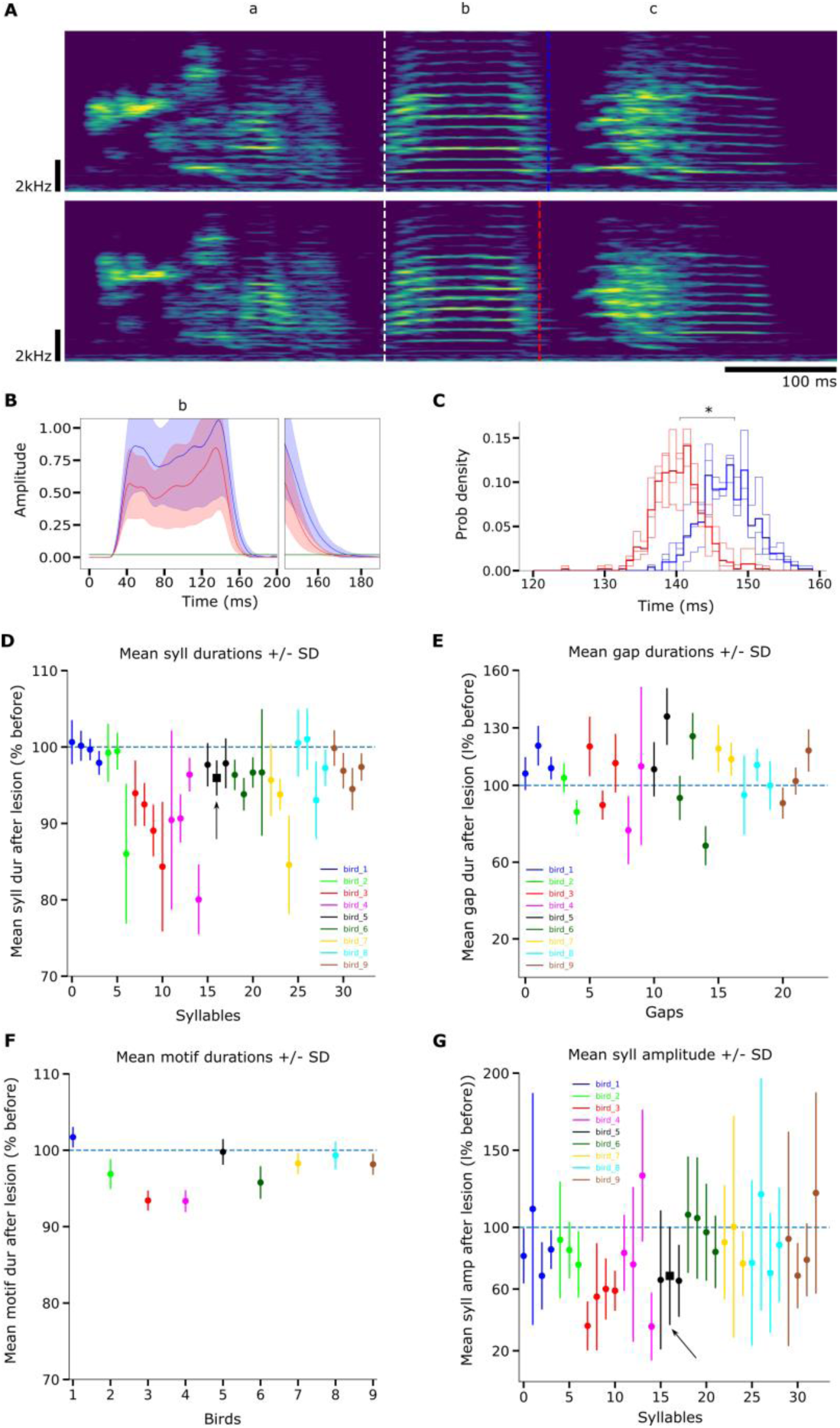
Bilateral lesions in the latDCN reduce syllable and motif duration and syllable amplitude, while inconsistently affecting gap duration. A: spectrogram of a song motif before (top) and after (bottom) bilateral lesion of the latDCN. The motif is made of three syllables (a, b and c). White dashed line: onset of syllable b, aligned for the two renditions. Blue and red dashed lines: offset of syllable b before and after lesion. B: Mean amplitude of syllable b across 3 consecutive days before (red) and after (blue) lesion (shaded region: standard deviation around the mean). In green, threshold used to measure syllable duration. C: Distribution of syllable b duration for three consecutive days before (blue) and after (red) the lesion. Dark blue and red: average distributions before and after lesion. D: Mean duration of all syllables before and after lesion in all birds (n=33 syllables for n=9 birds, vertical lines: +/- 1 SD, black arrow: example syllable from A-C). E: Mean duration of gaps before and after lesion (n=23 gaps for n=9 birds). F: Mean motif duration (n=9 birds) before and after lesion. G: Mean syllable amplitude before and after lesion (n=33 syllables for n=9 birds).

### Auditory inputs reach the cerebellum in anesthetized zebra finches

Given the role of the lateral cerebellum in syllable timing, we first examined the auditory and motor signals that could support internal models of vocal production. We began by probing auditory inputs in anesthetized finches. Prior studies in non-songbird species such as pigeons and owls indicate that auditory information is processed mainly in the posterior cerebellum, particularly lobules VII and VIII, and possibly lobules II–V (Gross, 1970; Whitlock, 1952). To assess the distribution and selectivity of auditory inputs in zebra finches, we recorded from large populations of neurons across multiple cerebellar lobules while presenting playbacks of the bird’s own song (BOS), reversed BOS, a conspecific song, and white noise. Lobules were numbered following standard conventions (Iwaniuk et al., 2006; Larsell, 1952; Voogd & Glickstein, 1998) (Suppl. Fig. 1F). Because the lateral deep cerebellar nucleus (latDCN) is implicated in the control of song timing, we focused on the cortical layers of lateral cerebellar lobules known to project to the latDCN across vertebrates (Voogd & Glickstein, 1998).

The cerebellar cortex contains many cell types, and extracellular recordings do not allow unambiguous identification of the recorded neurons. However, in mammals, action potential shape, spontaneous firing statistics, and the presence of complex spikes can help distinguish Purkinje cells from other neurons (Gao et al., 2012; Van Dijck et al., 2013). Purkinje cells are uniquely characterized by complex spikes followed by a pause in simple spike activity (Granit & Phillips, 1956; Ramirez & Stell, 2016). In our data, spike shape and spontaneous firing rate varied along a continuum, with spike width and firing rate providing the clearest separation among cells (Fig. 2E; Suppl. Fig. 2B). On 58 recording sites, complex and simple spikes were detected, and cross-correlation analyses confirmed that each complex spike was followed by a significant pause in simple spiking (Fig. 2E, green). These recordings are thus likely to include Purkinje neurons. Extracellularly recorded Purkinje cells do not always show detectable complex spikes, probably due to the electrode position. For sites showing both spike types with a significant post-complex spike pause, the firing rate of simple spikes was consistently higher than 12 Hz. Given the low firing rate of other cell types in mammals, we classified neurons into high firing rate (HF, putative Purkinje cells; red in Fig. 2E) and low firing rate (LF, putative interneurons or granule cells; blue in Fig. 2E) based on this threshold.

**Figure 2:**
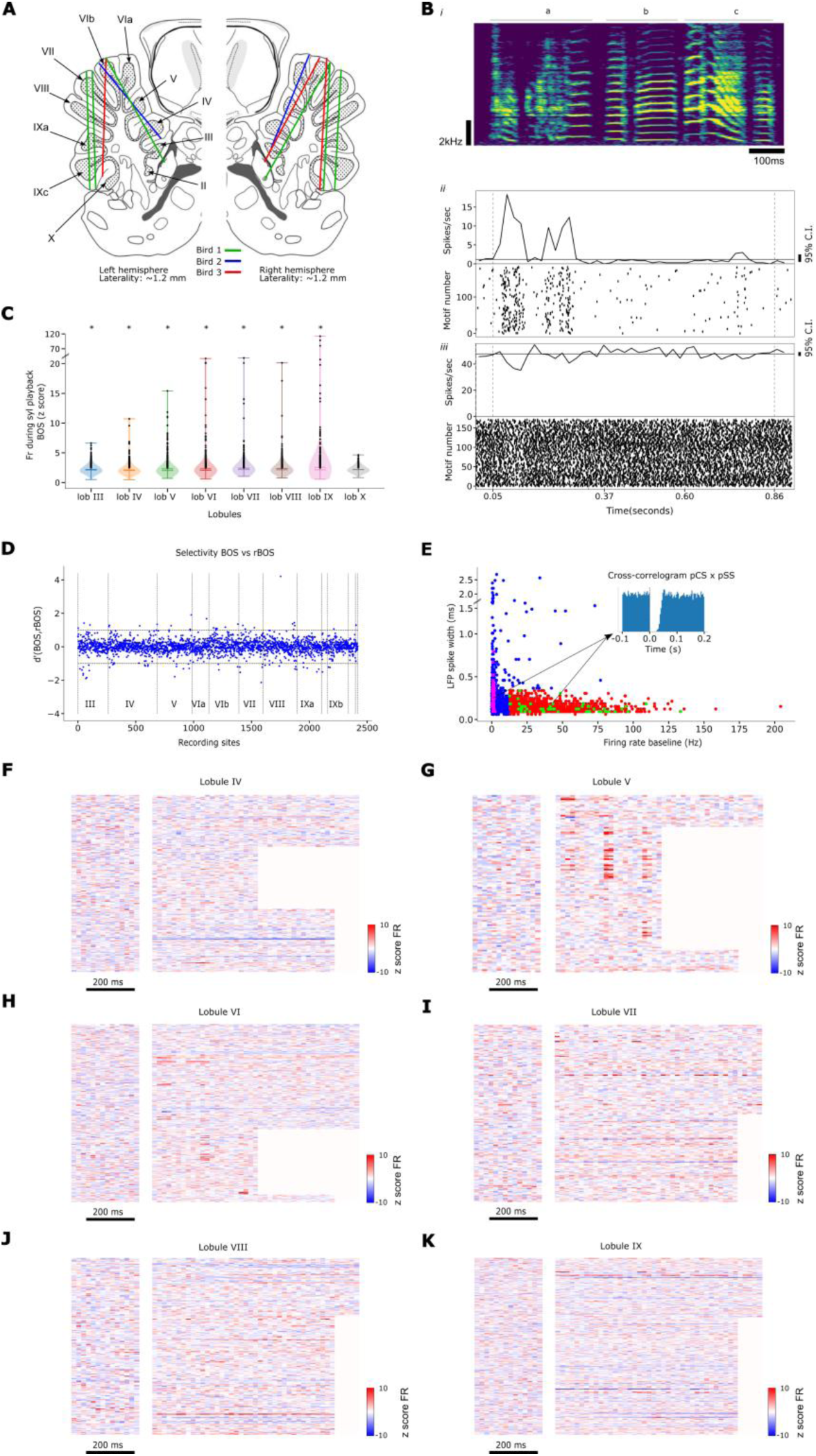
Auditory responses in the cerebellum of anaesthetized finches. (A) Neuropixel recording probe locations. (B) Top (i): example song spectrogram. Bottom: Example LF (ii) and HF (iii) units from lobule VI: raster plot (bottom) and PSTH of BOS playback (top, solid line: mean firing rate; solid horizontal line: baseline; vertical bar: 95% confidence interval; vertical dashed lines: motifs onsets/offsets). (C) Distribution of z-scored firing rates across lobules during BOS playback (n=3 birds). Black dots: units with z>2.5. Asterisks: lobules with above-chance proportions (bootstrap test, see Methods and Suppl. Fig. 3). Note that the y scale is piecewise linear. (D) BOS vs reverse BOS selectivity (d’ index per unit, see methods). Dashed lines: significance thresholds (d’>1, d’<-1). Units grouped by lobule (III–X, separated by vertical dashed lines). (E) Cell classification by baseline firing rate and spike width. Inset: cross-correlogram between complex spikes (CS, magenta) and simple spikes (SS, green). Blue: low frequency units, red: high frequency units. (F–K) Neuronal responses to BOS playback across lobules IV–IX. Each line: z-scored PSTH (20 ms bins) averaged across trials. Left: baseline. Right: BOS playback (duration = motif length, bird-specific). BOS motifs were presented at least 5 times and each recording contained at least 20 spikes to compute this average response. Only sites with |z|>2.5 are shown. Lobule IV: 221 recording sites shown. Lobule V: 118 recording sites shown. Lobule VI: 230 recording sites shown. Lobule VII: 179 recording sites shown. Lobule VIII: 202 recordings shown. Lobule IX: 321.

Both HF and LF neurons across lobules responded significantly to BOS playback (z-scored firing rate > 2.5; Fig. 2C). Except in lobule X, the proportion of responsive neurons was greater than expected by chance (Suppl. Fig. 3; Methods). In lobules V and VI, subsets of neurons showed robust, phasic, time-locked responses to BOS playback, as illustrated by individual examples (LF neuron, Fig. 2B,*ii*; HF neuron, Fig. 2B,*iii*) and population responses (Fig. 2G–H; Suppl. Table 1). By contrast, neurons in lobules VII, VIII, and IX tended to show sustained responses during BOS playback (Fig. 2I–K; Suppl. Table 1). Although BOS playback often evoked strong responses, reversed BOS, conspecific song, and even white noise also elicited activity changes (Suppl. Fig. 4). Response amplitudes and patterns (phasic vs. sustained) were comparable between BOS and white noise (Fig. 2F–K; Suppl. Fig. 4). Selectivity analysis confirmed that auditory responses were largely non-specific (d’ = 0.017 ± 0.39; Fig. 2D; Suppl. Fig. 5).

In summary, our recordings reveal that the lateral cerebellum receives broadly non-selective auditory input across nearly all lobules (Fig. 2F–K; Suppl. Fig. 6A,B), with particularly strong and phasic responses in lobules V and VI.

### Neuronal activity increases in the cerebellum during singing

While auditory responses in anesthetized zebra finches demonstrate cerebellar access to auditory information, recordings in awake, singing birds are necessary to reveal signals relevant to internal models and song timing. To this end, we implanted adult zebra finches with motorized microdrives and recorded cerebellar activity during spontaneous singing and passive auditory playback. To probe potential representations of expected vs. unexpected feedback, we introduced probabilistic Distorted Auditory Feedback (DAF, Fig. 3A–B). Auditory stimuli included BOS, reversed BOS, conspecific song, and white noise (Fig. 3C). We report the activity of 213 units (182 single units, 31 multi-units) from 15 birds. Recording sites, verified by electrode depth and post-hoc histology (lesion marks, Fig. 3D), spanned multiple lobules, with most in IV (n=4 birds), V (n=7), and VI (n=5) (Fig. 3E). Units were classified as HF (putative Purkinje neurons) or LF based on spontaneous firing rate and spike width, using thresholds derived from anesthetized recordings (Fig. 3F). Because Purkinje neurons fire faster in awake than anesthetized mammals, these thresholds are conservative (Sánchez-León et al., 2025). Alternative classification using inter-spike interval entropy (Van Dijck et al., 2013) did not improve separation (Suppl. Fig. 2A–B). One putative Purkinje cell displayed identifiable complex and simple spikes (Fig. 3H, Suppl. Fig. 2C–D). Both HF and LF neurons increased firing during singing compared to quiet wakefulness (HF: 68 ± 49 vs. 60 ± 50 Hz; LF: 11 ± 9 vs. 8 ± 5 Hz; paired t-tests, p<0.05; Fig. 3G). We next compared singing-related modulation to auditory-evoked responses across lobules to determine whether cerebellar neurons primarily encode motor commands or auditory signals.

**Figure 3:**
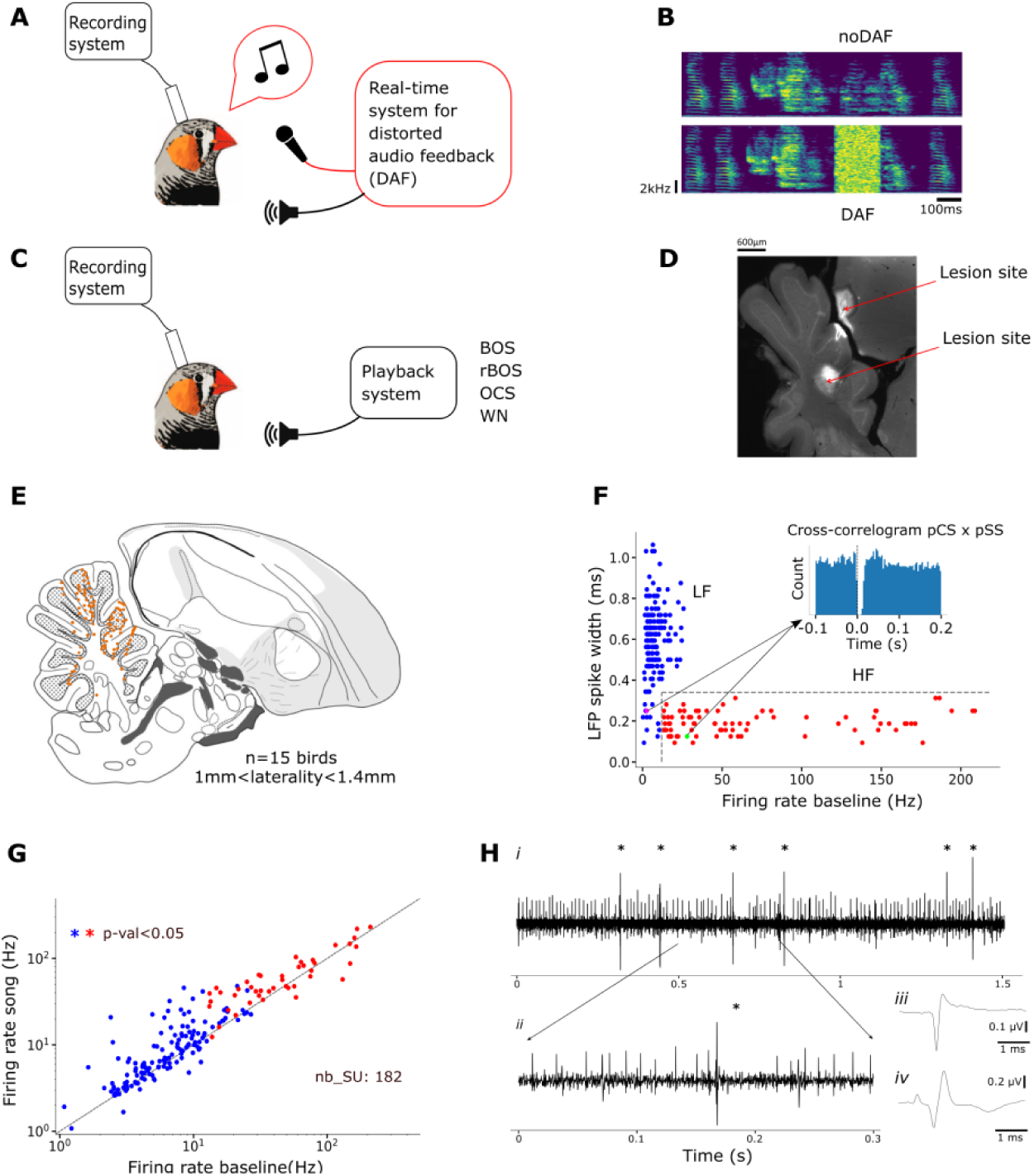
Neuronal activity in the cerebellum of awake, singing zebra finches. (A) Schematic of recordings during singing with or without probabilistic Distorted Auditory Feedback (DAF). A real-time system triggered white noise upon detection of a target syllable. (B) Example spectrograms of songs with and without DAF. (C) Schematic of passive playback recordings. Stimuli included BOS, reversed BOS (rBOS), conspecific song, and white noise. (D) Histological verification of electrode placement. Electrolytic lesions were made at two depths along the electrode track. (E) Recording site locations across lateral cerebellar lobules (n=15 birds; medio-lateral positions 1–1.4 mm). (F) Single-unit classification by baseline firing rate and spike width. Blue: LF units; red: HF units; green: putative simple spikes (SS); magenta: putative complex spikes (CS). Inset: cross-correlogram of SS and CS. (G) Firing rates were significantly higher during singing compared to quiet baseline for both HF (red) and LF (blue) units. (H) Example recording in lobule IV of an awake bird. (i) Raw trace showing putative CS (asterisks) and SS. (ii) Zoomed trace segment. (iii) Average SS waveform. (iv) Average CS waveform.

### Activity modulation during singing and song playback

As shown above, cerebellar neurons exhibited an overall increase in firing rate during singing compared to non-singing episodes. This elevation did not simply reflect a global arousal state but stemmed from strong time-locked modulations. Many neurons across several lobules displayed significant firing rate modulation during singing, illustrated here for an example LF neuron (Fig. 4A–C) and HF neuron (Fig. 4D,E) in lobule V. To assess the contribution of auditory feedback, we compared singing-related firing changes with responses to BOS playback during non-singing episodes. In some cases, such as the example LF neuron, responses during singing closely resembled playback-evoked activity (Fig. 4C). However, the relative strength of modulation during singing versus playback (z-score; see Methods) varied widely, even among neurons within the same lobule (48 LF and 12 HF neurons from 7 birds, Fig. 4G). In lobule V, LF neurons tended to show stronger modulation during playback than during singing (blue dots, p<0.05, related t-test), whereas HF neurons were more strongly modulated during singing than during playback (red dots, p<0.05, related t-test).

**Figure 4:**
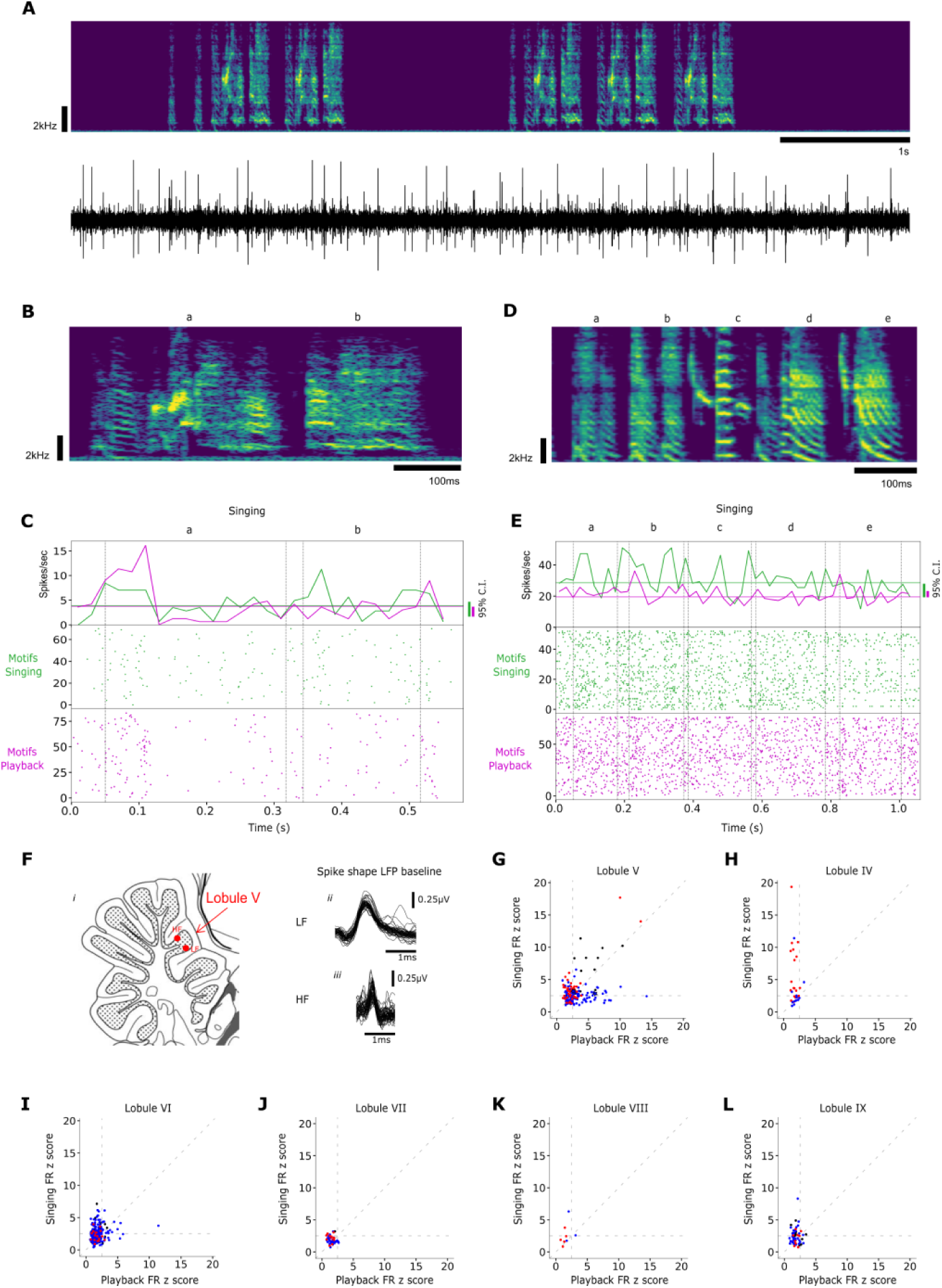
Singing-related and playback-evoked responses in the cerebellum, exemplified in lobule V. (A) Example LF neuron: spectrogram of a song bout (top) with raw recording trace (bottom). (B) Spectrogram of the song motif. (C) Raster plot (bottom) and PSTH (top) of the same LF neuron during singing (green) and BOS playback (magenta). Solid lines: baseline firing rate around singing (green) and playback (magenta). Vertical green and magenta bars: 95% confidence intervals for singing-related and playback-related PSTHs respectively. Vertical dashed lines: syllables onsets/offsets. This unit also responded to reversed BOS, conspecific song, and white noise (Suppl. Fig. 7). (D-E) Same as B–C for an example HF neuron in lobule V. (F) Recording sites (i) and spike shapes of the LF (ii) and HF (iii) neurons shown in C–E. (G–L) Comparison of peak z-scored activity during singing and BOS playback across lobules. Each point: one unit during one syllable. Red = HF cells; blue = LF cells; black = multi-units. Dashed lines: z = 2.5 and unity line.

Neurons with strong firing-rate modulation during singing were also observed in lobule IV (5 LF and 7 HF neurons from 4 birds, Fig. 4H), lobule VI (53 LF and 4 HF neurons from 5 birds, Fig. 4I), and lobule IX (12 LF and 3 HF neurons from 3 birds, Fig. 4L). In lobule VIII, only a small sample was obtained (2 LF and 1 HF neurons from 1 bird, Fig. 4K), preventing firm conclusions. Neurons in lobule VII showed little modulation during either singing or BOS playback (5 LF and 3 HF neurons from 2 birds, Fig. 4J). The relative proportions of neurons responding stronger during singing versus playback across lobules are summarized in Table 1. Notably, lobule IV neurons exhibited the strongest modulation during singing with minimal response to playback (Fig. 4H; see also below for detailed analysis). In most other lobules, the pattern resembled that of lobule V, with both singing- and playback-related modulation, substantial variability in relative strength across neurons, and a consistent bias for HF neurons to show stronger firing modulation during singing (significant in lobules IV, V, and VII; paired t-test, p < 0.05).

**Table 1:** Proportions of significant singing-related and BOS playback responses in the lobules IV, V, VI, VII, VIII and IX of the cerebellum.

| Nb (syllable, recording site)<br>significantly modulated during<br>($z > 2.5$) | Lobule<br>IV | Lobule<br>V | Lobule<br>VI | Lobule<br>VII | Lobule<br>VIII | Lobule<br>IX |
| --- | --- | --- | --- | --- | --- | --- |
| **singing only** | 48%<br>15/31 | 23%<br>36/160 | 32%<br>85/266 | 13%<br>6/45 | 25%<br>2/8 | 40%<br>21/53 |
| **singing and BOS playback** | 7%<br>2/31 | 29%<br>47/160 | 7%<br>18/266 | 0%<br>0/45 | 12%<br>1/8 | 6%<br>3/53 |
| **BOS playback only** | 0%<br>0/31 | 22%<br>35/160 | 4%<br>11/266 | 4%<br>2/45 | 0%<br>0/8 | 7%<br>4/53 |

As in anesthetized birds, cerebellar neurons in awake zebra finches did not show selective responses to BOS playback but instead responded non-selectively to BOS, conspecific songs, and reversed BOS (Suppl. Fig. 8). These responses were largely driven by sound onsets and offsets (Suppl. Fig. 7), contrasting sharply with the highly selective auditory responses observed in telencephalic auditory areas (Schneider & Woolley, 2013; Theunissen et al., 2004) or in forebrain song nuclei in anaesthetized birds (Konishi, 2004; Solis & Doupe, 1999).

During singing, cerebellar activity could in principle reflect auditory feedback, but it might also represent a corollary discharge of motor commands or an internal inverse model that links auditory and motor representations (Giret et al., 2014; Hanuschkin et al., 2013; Troyer & Doupe, 2000; Wolpert & Flanagan, 2001). These hypotheses make distinct predictions about the relative timing of auditory and singing-related activity. To test this, we compared PSTHs evoked during singing and BOS playback. In lobule V, for example, one HF neuron displayed highly similar PSTHs in both conditions (Fig. 5A). Cross-correlation analysis revealed a significant alignment with singing-related activity leading playback-evoked responses by ∼10 ms (Fig. 5B). Across the dataset, such alignment was found in 15% (11/71) of units in lobule V (Fig. 5D), but was rare or absent in other lobules (0–2 units, Suppl. Table 2). The average cross-correlation peak across these neurons (cc ≈ 0.5) was far below the level expected from pure auditory re-afference (cc = 1), yet significantly above chance, with a consistent ∼10 ms lead of singing-related over playback-evoked activity (Fig. 5D). Thus, while singing-related firing partially overlaps with playback responses, it cannot be reduced to an auditory re-afference. Instead, the temporal alignment of sensory- and motor-related signals in lobule V may rather reflect the implementation of internal models (Giret et al., 2014; Hahnloser & Ganguli, 2020; Wolpert & Kawato, 1998).

**Figure 5.**
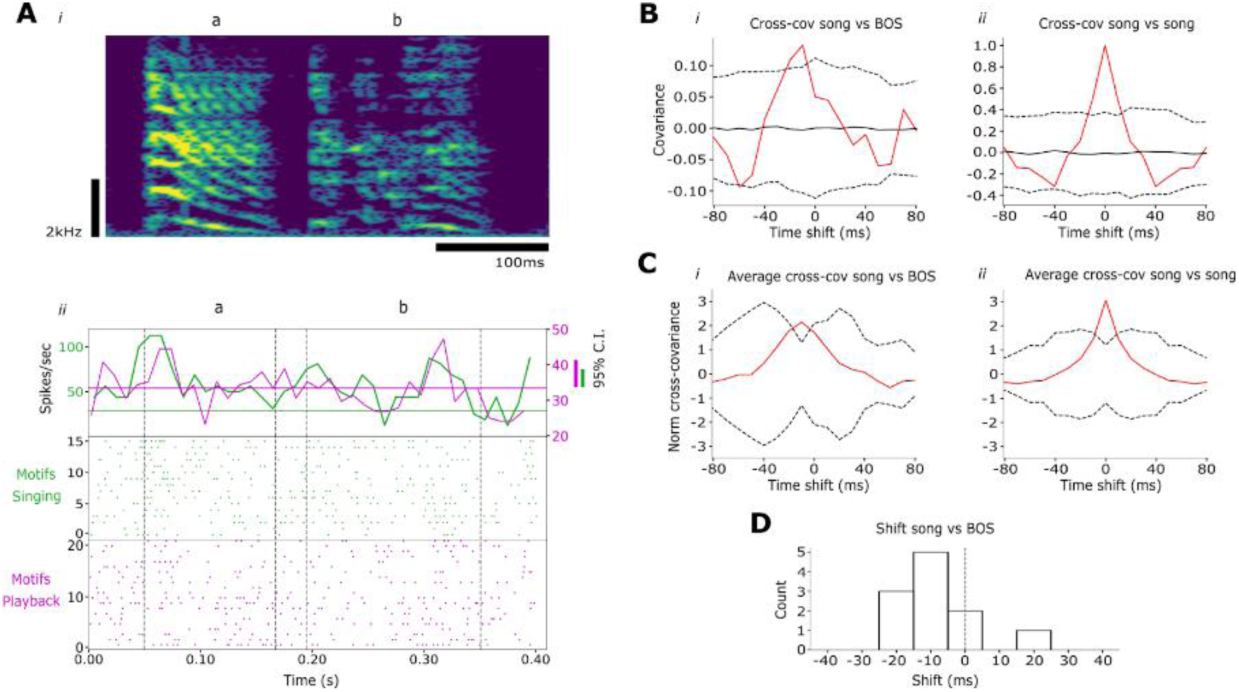
Alignment of singing- and playback-related activity in lobule V. A: Song spectrogram (top) ad example raster and PSTH during singing (green) and BOS playback (magenta). Vertical green and magenta bars on the side indicate 95% confidence limits for singing-related and playback respectively. Both conditions show partially aligned modulation, with increased firing after syllable a onset and before b offset. B: Cross-correlation between singing- and playback-related PSTHs shown in A (red). Shuffled-data mean (solid black) and ±2 SD interval (dotted black) are shown for comparison. Right: autocorrelation of the singing PSTH. C: Mean normalized cross-covariance between singing- and playback-related PSTHs across all units (red) with jackknife confidence interval (dotted black, see Methods). Right: corresponding normalized autocovariance of singing PSTHs. D: Distribution of peak latencies in cross-covariance for all recorded neurons showing a significant peak in singing and playback PSTHs covariance. Negative values indicate leading activity during singing compared to playback.

### Auditory feedback distortion does not affect singing-related activity

To test whether singing-related cerebellar activity reflects auditory feedback (re-afference), we recorded neuronal responses during spontaneous singing while birds were exposed to syllable-locked probabilistic Distorted Auditory Feedback (DAF; Fig. 6A, see Methods). If auditory feedback significantly contributed, DAF should alter singing-related modulation. However, while an example neuron strongly responded to white noise during passive playback (Fig. 6C), its activity during singing was unaffected by DAF (Fig. 6B,D). Across the population, many neurons exhibited robust responses to white noise during non-singing episodes (Fig. 6E), yet DAF failed to alter their firing during singing. Population analyses confirmed that white noise significantly increased firing rates during non-singing but not during singing (Fig. 6F–I). Although DAF may only partially overlap with the feedback experienced during singing, these results suggest that singing-related modulation in cerebellar neurons is not primarily driven by auditory feedback.

**Figure 6:**
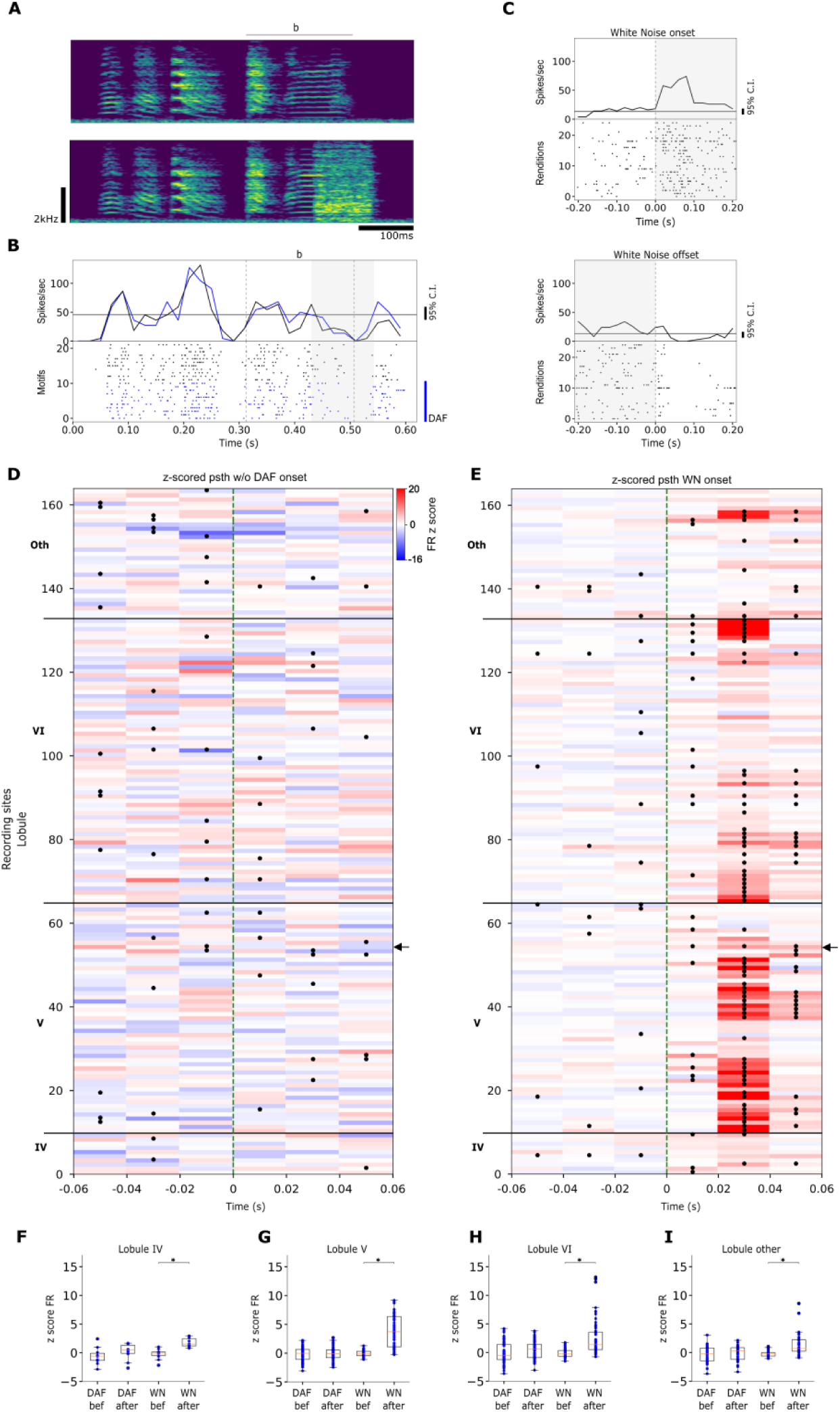
Effect of DAF on neuronal activity in lobule V. A: Spectrogram of a motif without (top) and with (bottom) DAF. B: PSTH and raster of an example unit during singing; blue dots and line indicate trials with DAF. Gray shade indicates the duration of the DAF. C: PSTH and raster of the same unit during white noise playback outside singing episodes (top: onset, bottom: offset). Gray shade indicates noise playback. D: Z-scored difference in firing rate between song renditions with vs without DAF across 164 sites in lobules IV–VI and others (20 ms bins). Green dashed line: DAF onset. Black asterisks: significant differences (t-test, p<0.05). Black arrow: example in B. E: Same as D for white noise playback onset outside singing episodes. Color code: z-scored firing relative to baseline. Black asterisks: significant changes (t-test, p<0.05, difference in firing rate between the bin of interest and a reference bin situated 100 ms before white noise playback onset). Black arrow: example in C. F–I: Z-scored average firing rates before and after noise onset during singing (DAF) vs playback (WN). Singing values represent the difference between renditions with vs without DAF. White noise significantly modulated firing during playback but not during singing (related t-tests: F–IV, -1.88 ns vs -6.12, p>0.05 vs p<0.001; G–V, -0.27 ns vs -9.88, p>0.05 vs p<0.001; H–VI, -1.76 ns vs -6.7, p>0.05 vs p<0.001; I–VII–X, -0.87 ns vs -4.17, p>0.05 vs p<0.001).

### Neuronal activity correlates with syllable duration

During singing, cerebellar internal models may either predict expected auditory feedback—leading neurons to modulate their firing in relation to song acoustics—or contribute to song timing, in which case firing should correlate with syllable duration. To test this, we assessed the correlations between the firing rates of cerebellar neurons during singing and both acoustic features (pitch, spectral entropy, amplitude, see (Sober et al., 2008)) and syllable duration (Suppl Fig. 9). To control for spurious correlations caused by slow firing fluctuations (Harris, 2021), we applied a detrending procedure (see Methods). Strikingly, a significantly larger-than-chance fraction of neurons showed correlations with syllable duration, both with detrending (29/186 cases, Suppl Fig. 10B) and without (55/186 cases, Suppl Fig. 11B). Bootstrap analyses confirmed the robustness of this effect in lobule V (Suppl Fig. 9D), across all neuron types (Suppl Fig. 10B) as well as within HF (Suppl Fig. 12B) and LF neurons (Suppl Fig. 13B). By contrast, correlations with acoustic features were rare, with only weak effects for pitch and spectral entropy in lobule IV HF neurons (Suppl Fig. 10E,I,M; Suppl Fig. 12E,I,M). Overall, we find little evidence that cerebellar firing tracks song acoustics, but consistent evidence for correlations with syllable duration, supporting a role for the cerebellum in internal models of song timing.

### Neurons in lobule IV tightly encode syllable boundaries

While neurons in lobules V, VI, VIII, and IX responded to BOS playback, lobule IV neurons showed strong modulation during singing but not during playback (Fig. 4H). Could this activity participate in the control of syllable duration by the cerebellum? Both LF and HF neurons increased their average firing rates during singing compared to silence (HF: 53 ± 39 vs 43 ± 40 spikes/s; LF: 15 ± 10 vs 9 ± 6 spikes/s; n=12 HF, 10 LF neurons, 4 birds). Firing was further modulated within song bouts, as illustrated by an example HF cell (Fig. 7A,D–E). Across all lobule IV recordings, 58% (18/31) of cases (one neuron during one syllable is single case) showed significant singing-related modulation (z > 2.5). This effect was stronger in HF neurons (z=5.87 ± 4.48) than LF neurons (z=2.91 ± 2.72), suggesting Purkinje cells as the main contributors. By contrast, BOS playback did not significantly modulate lobule IV activity (1/31 case above threshold, not different from shuffled baseline; p>0.05, Table 1).

**Figure 7:**
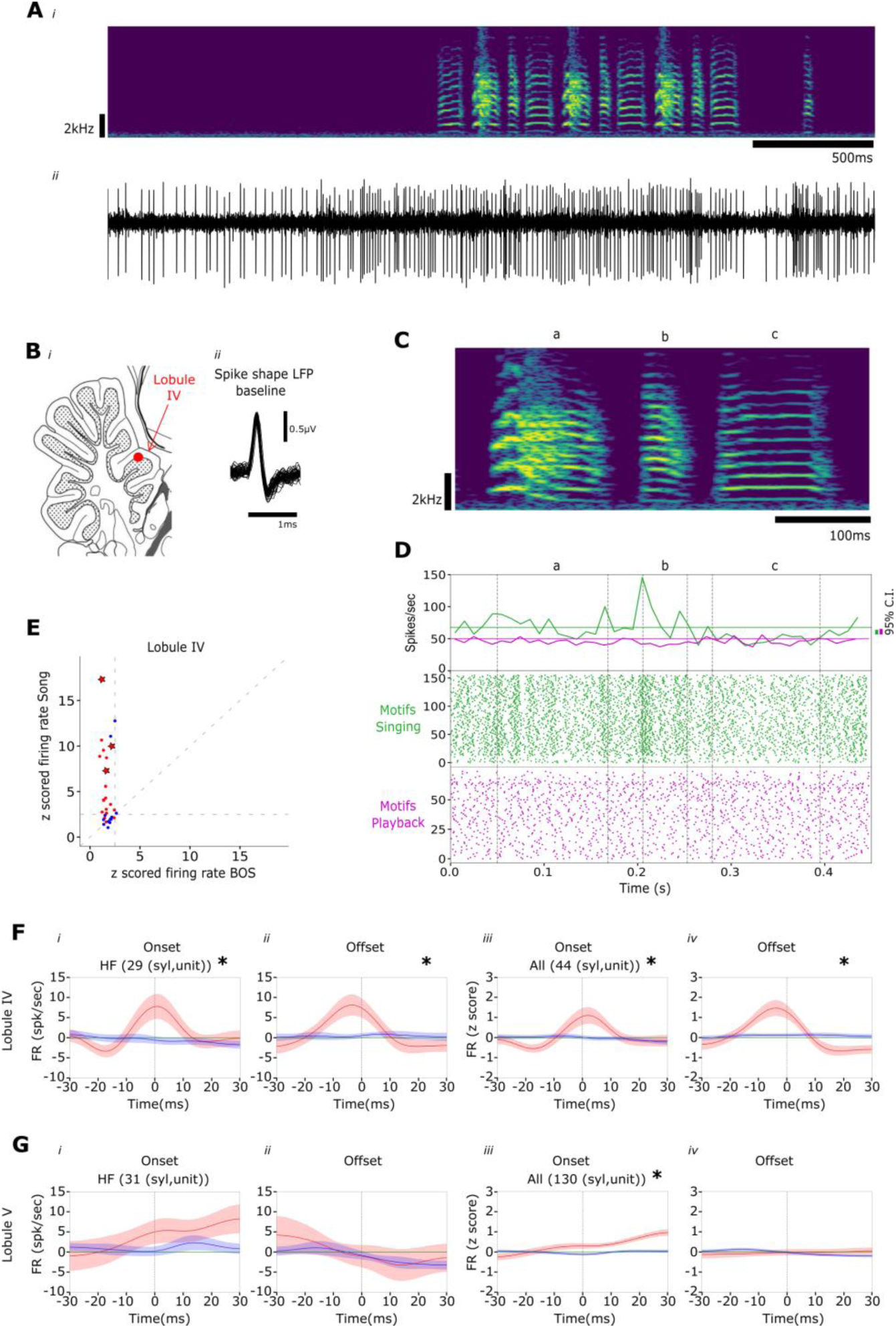
Singing-related activity is strongly modulated around syllable boundaries in lobule IV. A: Spectrogram of a song bout (top) and filtered LFP trace (bottom). B: Recording location (i) and spike shape of the recorded neuron (ii). C: Spectrogram of a song motif. D: Raster plot (bottom) and PSTH (top) of the same HF neuron during singing (green) and BOS playback (magenta). Solid lines: baseline firing rate around singing (green) and playback (magenta). Vertical green and magenta bars: 95% confidence intervals for singing-related and playback-related PSTHs respectively. Vertical dashed lines: syllables onsets/offsets. E: z-scored activity during singing vs BOS playback for all lobule IV neurons (n=3 birds). Each point: one unit during one syllable. Red = HF cells, blue = LF cells. Star-shaped points = example neuron from panel E during syllables a–c. F– G: Mean firing rate of HF units (i&ii) and all units (iii&iv) in lobule IV aligned to syllable onsets (i&iii) and offsets (ii&iv). Red: activity during spontaneous singing; blue: activity aligned to randomly sampled instants within motifs. The motif-averaged firing rate was subtracted; shading indicates ± SEM. Numbers in insets denote (syllable, unit) case number. Asterisks mark significant firing rate modulation at boundaries. G: Same analysis as F for lobule V.

Birds often display movements during song production. While undirected singing (the focus of our recordings) lacks the stereotyped courtship dance, it can still involve posture changes and head movements. Since cerebellar activity is generally influenced by body and head movements (Nguyen & Person, 2025), we tested whether the firing modulation observed during singing—but not BOS playback—could be explained by such movements. The strong, syllable-aligned firing modulation in lobule IV neurons (Fig. 4E), with reproducible peaks across motif renditions, already argues against this possibility, as body movements are not consistently aligned to song (Williams, 2001). To directly address this, we aligned neuronal activity to head and body movements tracked from video during singing (see Methods). Neuronal firing in lobule IV showed no significant modulation (Suppl Fig. 14A), indicating that the observed activity reflects vocal production rather than movement. As illustrated by the example neuron in Fig. 7D, many lobule IV neurons show firing modulations precisely aligned to syllable boundaries. To quantify this, we measured neuronal firing around syllable onsets and offsets (defined by song amplitude, see Methods). In lobule IV, the average firing rate across neurons exhibited sharp transients at both onsets and offsets (Fig. 7F), with half-widths of 16 ± 12 ms and 18 ± 9 ms, respectively. This temporal precision suggests that lobule IV neurons encode syllable boundaries, consistent with the reception of a motor efference copy during singing. By contrast, neurons in other lobules showed broader modulations around syllables (Fig. 7G; Suppl Fig. 15), incompatible with boundary encoding. Together, these results suggest that neurons in lobule IV encode syllable boundaries with high temporal precision, possibly representing expected syllable timing, and may thereby contribute to the cerebellar control of syllable duration.

## Discussion

We identified song-related signals across the lateral cerebellum in zebra finches. Auditory stimulation evoked strong but nonspecific onset responses in both anesthetized and awake birds, whereas neurons in many lobules exhibited singing-related firing modulations. The singing-related activity partially aligned with playback-evoked responses but remained insensitive to auditory feedback perturbations, ruling out a purely auditory reafference. Notably, neuronal activity in several lobules correlated with syllable duration rather than acoustic features, and firing in lobule IV was strongly modulated around syllable boundaries, together suggesting a role in the precise control of syllable timing.

The cerebellar song-related signals reported here likely arise from two distinct sources. First, nonspecific auditory responses may be mediated by brainstem, midbrain, or thalamic nuclei transmitting low-level auditory information(Hahnloser & Kotowicz, 2010). The lateral cerebellum receives direct input from cochlear nuclei (Wang et al., 1991) and indirect input via pontine nuclei from the inferior colliculus in mammals (Aitkin & Boyd, 1978; Brodal & Bjaalie, 1992; Kawamura, 1975). In birds, additional auditory input arrives from the auditory pallium through the medial spiriform nucleus (Gutiérrez-Ibáñez et al., 2018). Second, singing-related signals that differ from auditory reafference are more likely to reflect a corollary discharge of motor commands, as shown in rodents (Person, 2019) and predicted by internal model frameworks (Wolpert & Kawato, 1998). The premotor signals identified here may originate in the thalamic nucleus Uvaeformis (Uva), which conveys syllable onset signals (Danish et al., 2017; Moll et al., 2023), or from telencephalic song-control nuclei. However, firing patterns in lobule IV differ from those of Uva, HVC, or RA: HVC and RA neurons fire in sparse (Hahnloser et al., 2002) or dense patterns (Leonardo and Fee, 2005) across song, whereas Uva neurons increase activity before syllable onset and decrease it before offset (Danish et al., 2017). Thus, substantial processing likely transforms these inputs before reaching the cerebellum, possibly via Purkinje neurons that project to LatDCN. We excluded head and body movements as explanations for the observed firing modulations, but fine beak movements could contribute. Beak kinematics correlate with fundamental frequency but not with amplitude (Goller et al., 2004; Westneat et al., 1993), making them unlikely to account for syllable-boundary locking. They may, however, explain the small but significant correlation observed between cerebellar activity and syllable fundamental frequency (Suppl Fig. 10).

Our results show that lobule V activity during singing is partially aligned with auditory responses observed outside singing with a short time lag. The resemblance between auditory- and singing-related responses in this lobule is reminiscent of the “vocal mirroring” neurons described in HVC (Hamaguchi et al., 2014; Prather et al., 2008) and LMAN (Giret et al., 2014). Such mirroring has been linked to imitative associative learning (Hahnloser & Ganguli, 2020), and the observed alignment is consistent with an inverse internal model implemented in the cerebellum (Giret et al., 2014; Hanuschkin et al., 2013). At the same time, the robust onset-locked auditory responses to BOS and other stimuli suggest a role in imitation restricted to temporal song features such as syllable duration (Pidoux et al., 2018; Radic et al., 2024). Supporting this, lesions of the latDCN induced subtle but behaviorally relevant changes in syllable duration, as zebra finches discriminate timing differences with millisecond precision (Narula et al., 2018). Likewise, the sharp firing modulations in lobule IV (∼10 ms half-width) fall within the natural range of syllable duration variability (Glaze, 2006).

Neural dynamics in HVC tightly control the timing of song elements (Long et al., 2010; Long & Fee, 2008), and HVC participates in a larger loop involving RA, the brainstem, the thalamic nucleus Uva, and the forebrain nucleus NIf. Cerebellar output could contribute syllable timing information through interneurons projecting to the Dorsal Thalamic Zone (DTZ) (Nicholson et al., 2018; Pidoux et al., 2018). DTZ, in turn, projects both to Area X in the basal ganglia—which indirectly influences RA (Nottebohm et al., 1976) —and to the pallial nucleus MMAN, which provides input to HVC (Foster et al., 1997; Person et al., 2008; Vates et al., 1997; Williams et al., 2012). However, current evidence suggests that LatDCN output does not directly reach either the MMAN-projecting thalamic nucleus (*Nicholson PhD dissertation, 2018*) or Uva, implying that connections from LatDCN to HVC are more likely indirect. If mediated via DTZ or Uva, this pathway would involve at least three synapses, and further work is needed to clarify the precise connectivity.

The cerebellum plays a central role in movement timing and, more broadly, in sub-second temporal processing in the brain (Bareš et al., 2019; Sm et al., 1997; Tanaka et al., 2021). Clinical evidence shows that cerebellar lesions impair sensorimotor timing (Day et al., 1998; Flament & Hore, 1988; Ivry et al., 1988, 2002; Izawa et al., 2012). Specifically, the cerebellum regulates the duration of rapid, goal-directed and self-initiated movements (Kunimatsu et al., 2018; Miki et al., 2018), encodes the onset and offset of discrete gestures (Ma et al., 2022), and synchronizes actions with external cues (Okada et al., 2022). This temporal information is relayed to thalamo-cortical circuits for movement control (Dacre et al., 2021; Nashef et al., 2019).

In humans, cerebellar circuits likely contribute to multiple aspects of language production and processing (Ziegler, 2015). In songbirds, vocal learning circuits are thought to have evolved from ancestral motor learning pathways, a specialization process that may parallel the evolution of speech-related areas in the human laryngeal motor cortex (Davenport & Jarvis, 2023). The cerebellum itself is highly conserved across vertebrates, sharing common computational principles for motor learning (Raymond & Medina, 2018; Yopak et al., 2020). Thus, cerebellar circuits may have convergently adapted in both songbirds and humans to support vocal learning, producing functional similarities between birdsong and speech. Language tasks recruit lateral lobules VI and Crus I/II for sequencing motor commands, and lobules VIIb/VIII for phonological storage during verbal working memory (Mariën et al., 2014). Our findings in zebra finches show that lobules VI, VII, and VIII also carry song-related auditory and premotor activity. Consistently, lesions to the song-related basal ganglia in songbirds induce structural remodeling within the cerebello–thalamo–basal ganglia pathway, including lobules VIb and VII (Hamaide et al., 2018). This shared cerebellar topology across birds and humans may reflect the anatomical requirements for input–output access. Indeed, cerebellar circuits are organized into reproducible modular units spanning longitudinal and transverse axes, with localized DCN projections and functional segregation (Apps & Hawkes, 2009).

During speech, the human cerebellum likely supports premotor drive, temporal coordination, and sequencing of gestures across multiple timescales—from phonemes to phrases (Ackermann & Brendel, 2016; Bohland & Guenther, 2006; Ghosh et al., 2008)—with activity linked to speech rate (Riecker et al., 2006; Späth et al., 2022). This parallels the role of the avian cerebellum in controlling syllable and motif duration (Pidoux et al., 2018; Radic et al., 2024). In particular, the precise encoding of syllable boundaries in lobule IV, together with auditory–motor alignment in lobule V, suggests that cerebellar circuits implement internal models of vocal element duration. A comparable mechanism may underlie syllable timing control in human speech.

## Acknowledgments

We thank Nicolas Giret and Christelle Rochefort for their comments on the manuscript, Daniela Popa, Annemie Van der Linden, David di Gregorio and Clément Léna for thoughtful discussions and Remya Sankar for help with the experiments. We are greatly thankful to the technical staff of the Institut des Maladies Neurodegeneratives (IMN, CNRS UMR 5293) that provided support for this study. This work was supported by funding from the Agence National pour la Recherche (16-CE37-0020-01, 20-CE37-0023-02, 22-NEUC-0002-01), Fonds pour la Recherche Médicale (EQU202203014688) to AL, and from the Bordeaux International Graduate Program to EGZC.

## Materials and Methods

### Animals

All animals used in the experiments were adult male zebra finches (Taeniopygia guttata) aged beyond 90 days post-hatch. The birds came from our breeding facility or were bought from an external supplier (Oisellerie du Temple, L’Isle d’Abeau, France). The birds had continuous access to water and food and were housed under natural dark/light conditions.

### Real-time Distorted Auditory Feedback (DAF)

For the real-time DAF, we used the RTXI: Real-Time Experimental Interface (Patel, George, Dorval, White, Christini, & Butera, 2018). Briefly, a Digital Acquisition Card PCI-6221 is connected to the motherboard of a desktop computer when the RTXI is installed. The RTXI installation is documented on (www.rtxi.org) and consists in the installation of a Linux kernel that is then patched to receive real-time capabilities through the addition of the Xenomai framework. Once the desktop has booted into this new real-time kernel, the RTXI software is installed. To perform our real-time task consisting in audio feedback perturbation, we modified the default Data_Recorder part of the RTXI code similarly to (Skocik & Kozhevnikov, 2013). The code for the real-time audio feedback perturbation consists in computing every ms the correlation coefficient between a template spectrogram (the template is usually 30 to 50 ms long) and the spectrogram of the lastly recorded piece of audio signal (same duration as the template). We used RTXI to deliver white noise at a specific time in the motif, usually at the onset of a given syllable. We set the amplitude of DAF at the same level as the amplitude of the targeted syllable.

### Surgery

Before surgery, birds were first food-deprived for 30-40 min, and an analgesic was administered just before starting the surgery (meloxicam, 5 mg/kg). The anesthesia was then induced with a mixture of air and 4% isoflurane for 1 min then maintained for several minutes at 0.7-1.5% isoflurane to verify that the breathing rhythm is stable. Birds were then moved to the stereotaxic apparatus and maintained under anaesthesia with 0.7-1.5% isoflurane. Lidocaïne (16.22 mg/mL) was applied under the skin before opening the scalp. Small craniotomies were made above the midline reference point, the bifurcation of the mid-sagittal sinus, and above the structures of interest. Stereotaxic zero in the antero-posterior and mediolateral axis was determined by the sinus junction. Usually the cerebellum was accessed using a head angle of 50°. The stereotaxic coordinates used for each brain structure are summed up in Table 1.

**Table 1:**
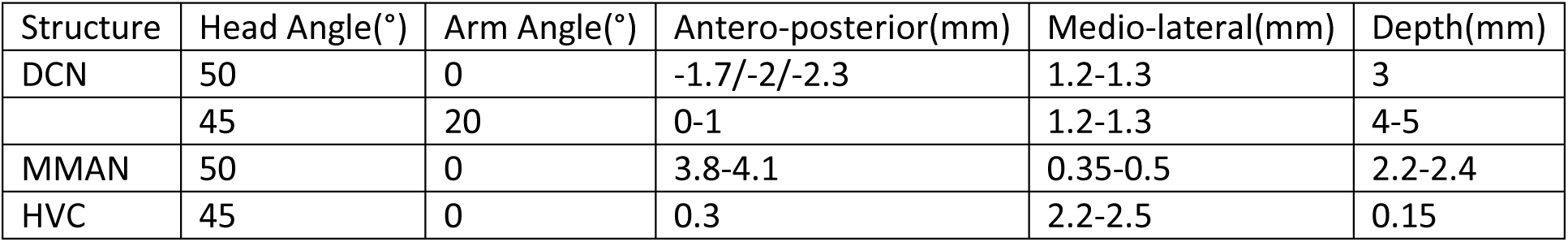
Coordinates for stereotaxic surgery.

| Structure | Head Angle(°) | Arm Angle(°) | Antero-posterior(mm) | Medio-lateral(mm) | Depth(mm) |
| --- | --- | --- | --- | --- | --- |
| DCN | 50 | 0 | -1.7/-2/-2.3 | 1.2-1.3 | 3 |
|  | 45 | 20 | 0-1 | 1.2-1.3 | 4-5 |
| MMAN | 50 | 0 | 3.8-4.1 | 0.35-0.5 | 2.2-2.4 |
| HVC | 45 | 0 | 0.3 | 2.2-2.5 | 0.15 |

### Lesions

Lesions were performed in the DCN of zebra finches. We targeted the most lateral DCN, analogous to the dentate nucleus in mammals. In a group of birds (n = 2), a partial electrolytic lesion was performed in the lateral deep cerebellar nucleus by passing 25 µA during 60-90 s through a tungsten electrode. Lesions were made at three points on the antero-posterior axis (see the stereotaxic coordinates in Table 1, DCN coordinates, second row). In a second experimental group (n = 7), chemical partial lesion was performed by pressure injecting ibotenic acid (10µg/µl) (Picospritzer III, www.parker.com/precisionfluidics), with a rate of 100 nL/min. We also performed injections at three locations along the antero-posterior axis (see Table 1, DCN coordinates) injecting 80-150nL per point. Following surgery, the behaviour of birds was closely monitored for a few days to ensure proper recovery. All birds underwent temporary motor deficits (postural and balance troubles). Most birds recovered quickly and were perching and feeding normally 48-72 hr after surgery. Singing usually resumed after 72 hr. Some birds (n=5) died 24-72 hours after surgery. In an attempt to reduce the mortality, we decreased the injected volume of ibotenic acid to 80 nL per injection site for several birds. We report in this study the duration of syllables, gaps and motifs before and after the surgery (>5 days post-surgery for full recovery) for the birds undergoing latDCN lesions or sham surgery.

### Electrophysiology

#### Freely moving

The electrophysiological recordings were performed using either tungsten electrodes (mostly 10MOhm but sometimes also 20MOhm) from FHC or quartz tetrodes (1-2MOhm type D or 4MOhm type A) from Thomas Recording. The electrodes were mounted on an electric motor (Faulhaber type: 0308A003B 125:1 S3) which allowed the insertion of the electrodes into the brain tissue remotely while the bird was singing (similar to (Fee & Leonardo, 2001)). The motor was remotely controlled with an RP Metrix MC2 controller (www.rpmetrix.com). Typically the electrodes were lowered through flexible guidance tubes with a speed of 10µm/sec by steps of 5-10 µm until a neuron was found. The neurons were recorded until the bird sang 100-200 songs. If the bird was not singing for long periods, we played the audio stimulation to the bird, including playbacks of BOS, reversed BOS a conspecific song and a white noise stimulus. The different stimuli were organized in a random sequence with 1-2 seconds between stimuli. If the bird resumed spontaneous singing, we stopped the auditory stimulations. The recording was done using 4 channels (either 4 tungsten electrodes or the 4 channels from the tetrode). The wires carrying the electrophysiological signals were soldered to a PCB attached to the motorised drive and then routed via an omnetics connector to a Neuralynx designed head-stage. The head-stage pre-amplified and digitalised the signals. The head-stage was attached via a flex cable to a Neuralynx Saturn commutator or directly to an Imetronic commutator (https://www.imetronic.com/). Finally the signals were processed by the Digital Lynx SX recording system and visualised on Neuralynx Cheetah software. The electrophysiology signals were sampled at 32000 Hz and band-pass filtered with cut off frequencies at 0.1 and 9000 Hz. The spikes signal was obtained by band pass filtering the signal with cut off frequencies at 200 and 3000 Hz. The audio signal was amplified via the Steinberg UR44 amplifier, digitalised and band pass filtered by the Digital Lynx SX with cut off frequencies at 0.1 and 9000 Hz. During recording, birds were video recorded with a web-cam (WideCam F100) and the videos compressed (MPEG2+MPGA). Some implanted drive featured a small lateral screw capable of slightly displacing the flexible guidance tubes. This allowed us to change at the end of the day the anterior-posterior location of the insertion place of the electrodes for the next day. To confirm the location of the electrodes, we either performed electrical lesions at different depth in case of tungsten electrodes or covered the electrode with a dye (Dil stain) before the implant in case of quartz tetrodes. For electrical lesions, after all the possible sites were recorded, we anesthetised the bird on a surgery table and we passed a DC current of 25 µA for 30-60 sec at different depth through one electrode. The location of the lesions or of the dye was then confirmed histologically (Fig 3D). The first two lobules (I and II) are very small and difficult to access in the lateral cerebellum and we did not record from them.

#### Anesthetized

The recordings in anesthetised birds subjected to audio stimuli were done using the Neuropixels probes. The probes were briefly introduced in a solution of DiI before being inserted in the cerebellum at a speed of 5-6 µm/s. The recordings started 30 to 40 min after the end of the insertion of the probes in order to allow the tissue to stabilise. After the recordings, the birds were euthanized with an isoflurane overdose; the brains were extracted and put in PFA 4% for 24 hours.

### Histology

After the electrophysiology recordings: Birds were sacrificed with a lethal intraperitoneal injection of Exagon (pentobarbital sodique, 400 mg/mL), perfused intracardially with PBS 0.01M sometimes followed by 4% paraformaldehyde as fixative. The brain was removed, always post-fixed in 4% for 24 hr, and cryoprotected in 30% sucrose. We then cut 50µm thick sections in the parasagittal or horizontal (for cerebellum) plane with a freezing microtome. Slices were mounted with Mowiol (Sigma Aldrich) and observed under an epifluorescence microscope. Images were analysed using ImageJ software (Rasband WS, NIH, Bethesda, Maryland, USA).

To quantify the extent of the lesions, birds were perfused and brains sliced (50µm thick) in a horizontal plane as described above. The slices were Nissl stained to measure the efficacy of the lesion by counting the number of neurons in the lateral DCN using the Mercator software (Explora Nova, La Rochelle, France).

### Data analysis

#### Spike sorting

For the recordings done with tungsten electrodes, spike sorting was performed with Spike2 software (CED, UK) using principal components analysis of spike waveforms. Spike train analysis was then performed using Python. For the recordings done with Neuropixel electrodes, spike sorting was performed using Kilosort within SpikeInterface. Electrophysiologically recorded units with SNR>5 and with less than 2.5% of Inter spike intervals below 1ms were classified as Single Units (SU). The rest were classified as Multi Units (MU). In anesthetised recordings, the single units with a firing rate below 3Hz were classified as putative Complex Spikes (CS) by visually inspecting the spike shape. For each putative complex spike, all the single units located within a distance of 300µm and having a firing rate above 12Hz were considered as potential putative Simple Spikes (SS). A cross-correlogram was built for each pair of putative complex spike and potential putative simple spike. The pairs featuring a significant pause in the cross-correlogram were considered as associated putative complex spikes and simple spikes. The pause in the cross-correlogram was considered significant if it appeared within 50 ms after time 0 and had at least two consecutive bins with a z-score below 2units of standard deviation below the average z-score of the cross-correlogram. The largest spike width of the putative simple spikes in the anesthetised recordings together with the lowest firing rate of a simple spike found in the literature (Dugué, Tihy, Gourévitch, & Léna, 2017) determined the borders of the High Firing (HF) units. The single units that were not classified as high firing units were classified as Low Firing (LF) units.

#### Statistics

We calculated peri-stimulus time histograms (PSTH) of recorded neurons with a bin of 10 or 20 ms. We calculated the mean and the 95% confidence interval of the firing rate over a baseline period. For song related PSTH the baseline period was a silent and quiet period (bird not singing and not moving). For audio stimulation related PSTH, the baseline period was made of several intervals in-between the audio stimuli. For some analysis, when stated explicitly in the legend of the figures, the baseline was set within the song motif or within the audio stimulus.

The PSTH around syllable boundaries was calculated only for syllables with a maximal intrasyllable z-scored firing rate above a threshold of 2.5. The mean and standard deviation of the firing rate for the z-score was measured during the motif rendition. The bins used for the PSTH were set to 10 ms and the PSTH were smoothed with a gaussian kernel of 5 ms and averaged before plotting. For a given syllable and cell type, random time points were distributed inside the motif to compute the shuffled PSTH. For a given lobule and cell type (HF or LF) the mean PSTH was computed and plotted with +/- sem. To test the significance of the firing rate at syllable boundaries, we performed a related t-test (using scipy.stats.ttest_rel()) between the measures of the firing rate at syllable boundaries and the firing rate and randomly distributed times during the motifs. The level of significance was chosen to be p=0.05.

To test whether the firing rates of a unit during clean renditions of a motif were different from the noisy renditions (renditions with DAF), we performed a related t-test (using scipy.stats.ttest_rel()) between the clean and noisy PSTHs. A PSTH was computed for each motif rendition and the same number of clean PSTHs and noisy PSTHs was used for the test. The bin size for the PSTHs was either 10 ms or 20 ms.

To test whether the firing rates of a unit during white noise playback onset were significant, we performed a related t-test (using scipy.stats.ttest_rel()) between PSTHs of the unit inside a 80ms window around white noise playback onset and the PSTH of the unit outside that window during the inter-stimulus interval. The bin size for the PSTHs was either 10ms or 20 ms.

To test whether the lesion of the DCN significantly affected the duration of song elements, we first tested the hypothesis that the data was drawn from a normal distribution using a Shapiro test (scipy.stats.shapiro()). If the hypothesis of normality of the data was kept (p-val>0.05), we performed a related t-test (using scipy.stats.ttest_rel()) between the mean durations of the elements before and after lesion. If the hypothesis of normality was rejected (p-val<0.05), we tested the hypothesis that the log transformed data was drawn from a normal distribution using the Shapiro test. If the hypothesis of normality of log transformed data was kept (p-val>0.05), we performed a related t-test on the log transformed mean durations of song elements before and after lesion. If log transformed data are used, this is explicitly specified in the text. For the figure 1,F, one out of nine birds had a non-consistent motif sequence consisting either of the syllables b, c, d or e, c, d. with syllables ‘b’ and ‘e’ having similarities in their spectrogram but different durations. The lesion significantly reduces the mean duration of the motifs independently of the motif sequence chosen for that bird (p-val<0.05 in both cases).

For the statistical significance we used the value of 0.05 as threshold for the p-value. For the supplementary figures 10,11,12,13, we used a Bonferroni correction or multiple comparisons. The threshold for the p-value in that case was chosen as 0.05/16.

#### Videos

During electrophysiological recordings, birds were recorded using webcams. We used the openCV python library to analyse the videos (https://docs.opencv.org/4.x/index.html). The videos were converted in grayscale format before being processed. The movement of the birds was quantified by processing the sequence of the difference of consecutive video frames. For each frame, the difference between two consecutive frames was smoothed using a two-dimensional gaussian kernel. A movement onset was detected if at least one pixel in the smoothed difference of consecutive frames exceeded a threshold. The threshold was determined manually after visual inspection of the video of interest. The timings consecutive movement onsets had to be spaced by at least 400 ms.

#### Songs sorting

We performed song segmentation into syllables using the supervised machine learning tool hybrid-vocal-classifier or hvc (Nicholson D., 2018). Briefly, raw audio signals were band pass filtered with cut off frequencies at 500 and 8000 Hz. The resulting signal was squared and low-pass filtered with a box car window of 10ms. This gave the envelope of the song signal. The song envelope was segmented by selecting the segments of the envelope that are above a manually selected threshold. The segments of around 100 to 200 motifs were manually labelled and used to train the hvc software (that used the SVM algorithm). The subsequent songs of the same bird were then automatically segmented by the so trained software. The output of the segmentation is a list of onset and offset times of each segment as well as the label of the segment that can be a syllable or a noise. For some birds it was not possible to find a threshold value allowing to segment all the syllables perfectly. The quality of the segmentation was verified using custom scripts and erroneously labelled syllables were removed from the analysis. A syllable was considered correctly labelled if the label was inside the sequence of labels building the motif. In addition, the duration of the syllable had to lie with +/-3 standard deviations around the median duration value; else the syllable was visually inspected and manually removed from the data set if it was not matching the expected syllable type.

#### Song-related activity

To build the psth during singing we first performed a piecewise linear Dynamical Time Warping (DTW) on the neuronal recordings. The mean of each syllable and gap was computed and the neuronal activity during each rendition of the syllable or gap was time warped (stretched or squeezed) to the mean duration.

For measuring the significance of the alignment between the singing evoked PSTH and the playback (BOS) evoked PSTH, we computed the cross-covariance between the PSTHs. We also computed the cross-covariance between the singing evoked PSTH and the shuffled playback evoked PSTH. The recording sites for which the cross-covariance exceeded the mean shuffled cross-covariance +/- 2 std for at least two consecutive points in time were considered featuring a significant alignment of motor and sensory evoked PSTHs (see figure 5B). For quantifying the significance of the alignments at the population level, we considered all the recording sites in lobule V featuring a significant alignment of song evoked and playback evoked PSTHs and computed the average normalized cross-covariance. The normalization of the cross-covariance was done by dividing the covariance by the jackknife estimated standard deviation (as in Giret et al., 2014)

To correlate the neuronal activity with the spectral or temporal features of the song, we computed the deviation of the firing rate from the mean firing rate during a time window. We then computed the deviation from the mean of the spectral or temporal feature of the syllable during another time window. The time window for the song feature was defined by the onset and offset of a syllable sub-tone or the whole syllable. The time window for the neuronal activity was defined by the onset and offset of a syllable in case of the syllable duration. The time window for the neuronal activity contained an additional 50ms wide premotor window before the syllable onset for the pitch, amplitude and spectral entropy.

For the correlation analysis, the firing rate of the units was first corrected for drifting activity using a detrending procedure. Detrending the firing rate of a unit consisted in plotting the cumulative number of spikes versus the spike times and manually selecting the breaking points of the curve. Then the instantaneous firing rate of the unit was computed by smoothing the spike train with a 20ms Gaussian kernel. Finally the firing rate was detrended within each region defined by the breaking points using the python detrending function signal.detrend(). The detrended firing rate was used to measure the correlation between the firing rate and the temporal and spectral features of the syllables.

The bootstrap analysis consisted in shuffling the correspondence between song renditions and neuronal activity within each recording site for all the recording sites and recomputing the correlations. The shuffling was done 1000 times which gave a distribution of the number of significant correlations. This distribution is shown in blue in supplementary figure 9D and supplementary figures 10,11,12,13. The number of significant correlations found in the real, non-shuffled data set is shown in red in the same figures.

The selectivity of the units to different audio stimuli presented in figure 2D and supplementary figure 5 was measured using the d’ metric proposed in (Solis & Doupe, 1997). This metric measures the similarity between the firing rates during a pair of auditory stimuli.

To compensate for the reduction in syllable amplitude due to the lesion, we measured for each syllable the average amplitude before and after lesion and computed a change ratio. We rescaled the amplitude of the syllables after lesion using the change ratio. We used the rescaled amplitude of the syllables to deduce the new values of syllable onsets and offsets as crossing points of the rescaled amplitude with the same threshold used without compensation. We then deduced the new syllable durations that we compared with the durations without compensation. The compensation was performed either only in case of a decrease in the amplitude after the lesion or in all cases. In both cases, the duration of the syllables significantly decreased after the lesion.

## Supplementary figures

**Supplementary figure 1:**
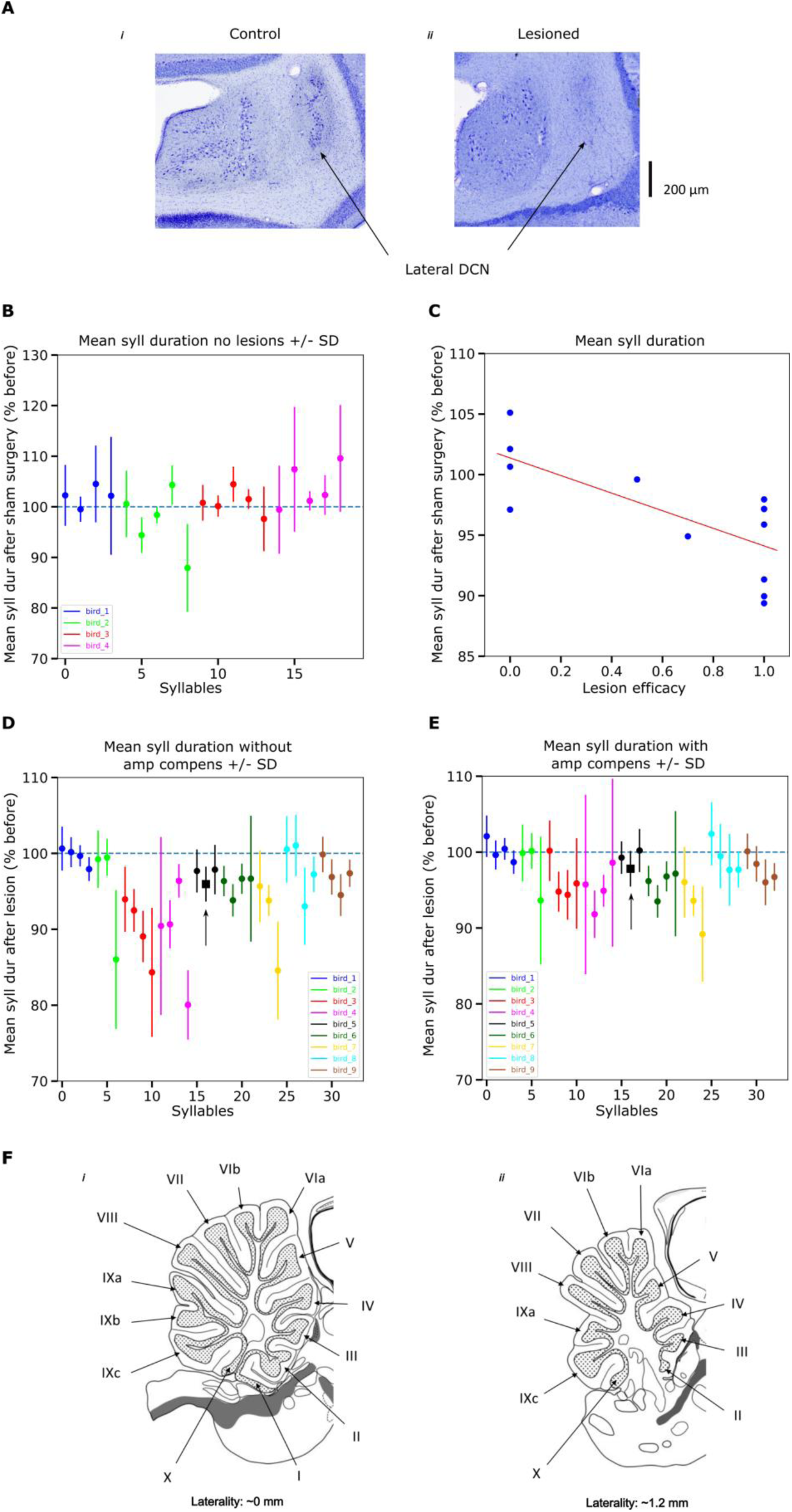
A: Nissl stained coronal slices of an example control bird (*i*) and a bird with lesioned lateral DCN (*ii*). B: Effect of sham lesions on control birds. The sham lesion didn’t significantly reduce the duration of syllables (t(19) = 0.79, p>0.05, related t-test on log transformed durations (see Methods)). C: Mean syllable duration as a function of the lesion efficacy. Each blue dot represents a bird. D: Same as Figure1D. E: Same as Figure1D but with compensation in reduction of syllable amplitudes (t(33) = -5, p<0.001, related t-test on log transformed durations). F: Numbering of the cerebellar lobules at the level of the sagittal symmetry plane (*i*) and at the level of the lateral cerebellum (*ii*). The laterality corresponds to the laterality of the lateral deep cerebellar nuclei.

**Supplementary figure 2:**
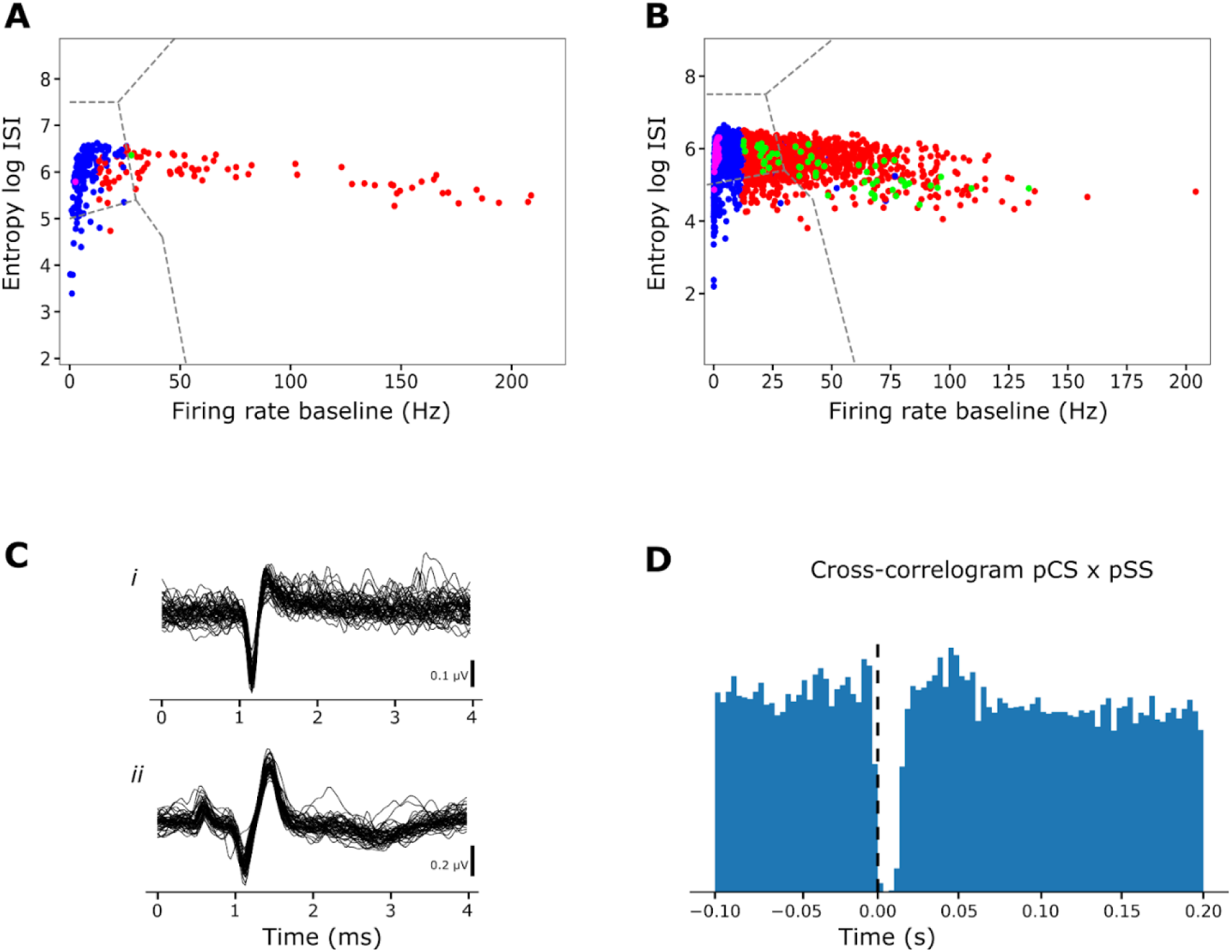
A: Classification of SU in awake birds according to the log of the ISI and firing rate during baseline as in (Van Dijck, et al., 2013). B: Same as in A but in anesthetised recordings. The colour codes are the same as in figure 3. C,*i*: Superimposed LFP traces of putative Simple Spikes from the recording shown in figure 3,I. A,*ii*: Superimposed LFP traces of putative Complex Spikes from the same recording. D: Cross-correlogram between putative Complex Spikes and Simple Spikes showing a pause in Simple Spike firing straight after the occurrence of a Complex Spike.

**Supplementary figure 3:**
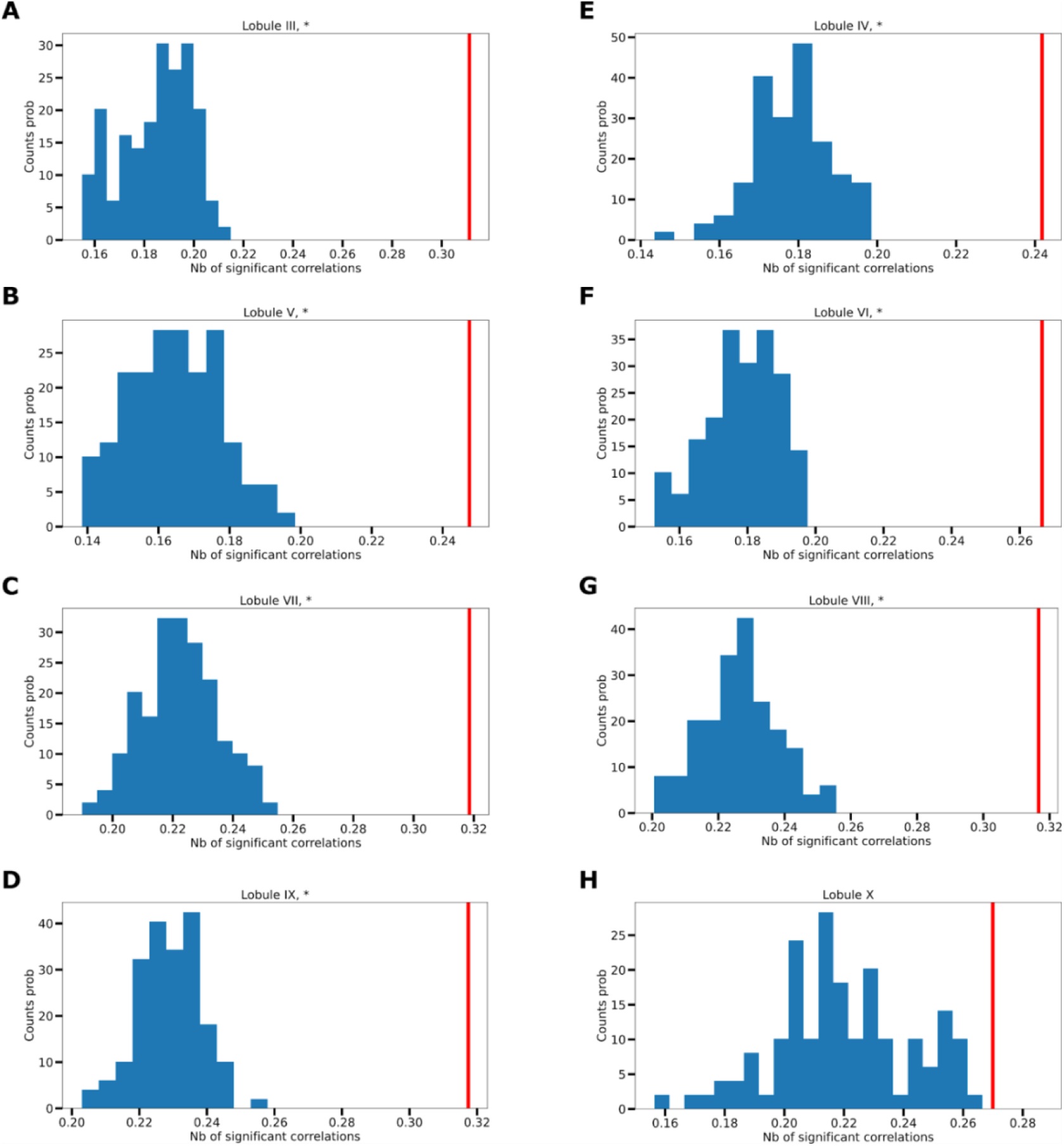
Significance of the responses to BOS in anesthetised birds. A: Histogram of bootstrap distributions of the proportion of cases (recording site x syllable) with z-score above 2.5 recorded in lobule III. The bootstrap was performed by measuring the proportion of z-scored firing rates above 3 during randomly distributed intervals outside the BOS stimuli. The procedure was repeated 100 times. The vertical red line represents the proportion shown in figure 2. The significance of the proportions from figure 2 is indicated by an asterisk. B,C,D,E,F,G,H: same as A but for the lobules IV, V, VI, VII, VIII, IX, X.

**Supplementary figure 4:**
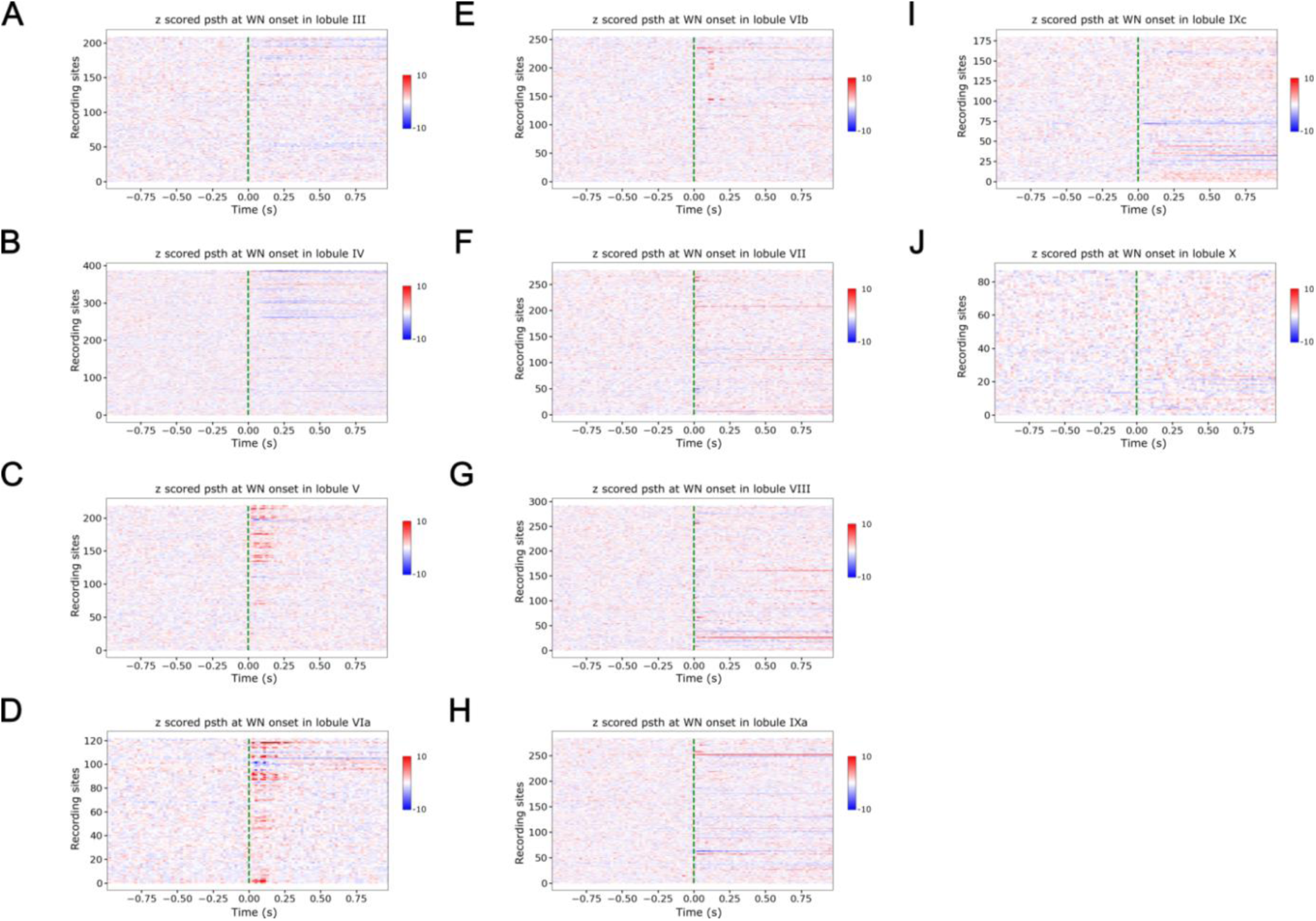
Response of the neuronal activity in cerebellar lobules to the onset of white noise. A: Each line represents the colour coded z-scored psth of a recording site in the lobule III. Only the recording sites are selected for which z>2.5 or z<-2.5 after noise onset. The green vertical dashed line represents the white noise onset. The bin for the psth is 20 ms. The colour code bar is on the right. B,C,D,E,F,G,H,I,J: same as A but for the lobules IV, V, VIa, VIb, VII, VIII, IXa, IXc and X.

**Supplementary figure 5:**
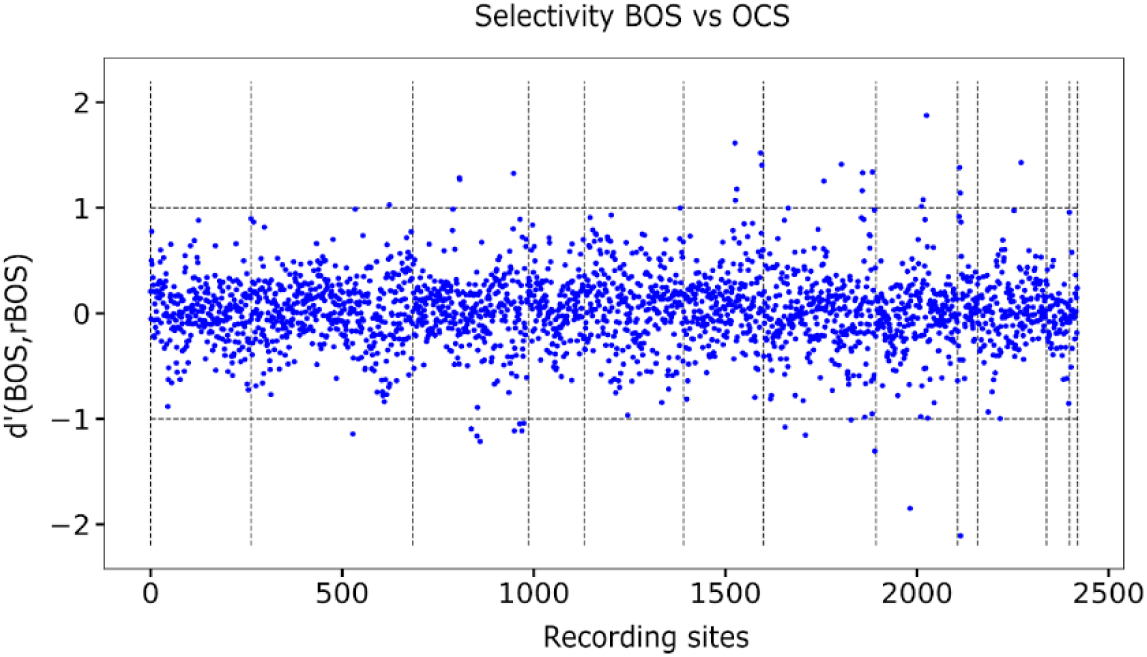
Selectivity of cerebellar neurons in anesthetised recordings. Each dot represents the selectivity of the neuronal activity of the recording sites to BOS vs conspecific song stimuli measured by d’. The horizontal dashed lines at d’=1/-1 represent the level of significance. d’>1 means the neuron is more selective to BOS that reverse BOS or conspecific song. d’<-1 means the neuron is more selective to conspecific song or reverse BOS than to BOS. The recording sites are sorted by lobule: lobule III, IV, V, VIa, VIb, VII, VIII, IXa, IXb, IXc, X, outside cb. The selectivity d’ was measured as in (Solis & Doupe, 1997) and is a normalised difference of the mean firing rate during the two types of audio stimuli.

**Supplementary figure 6:**
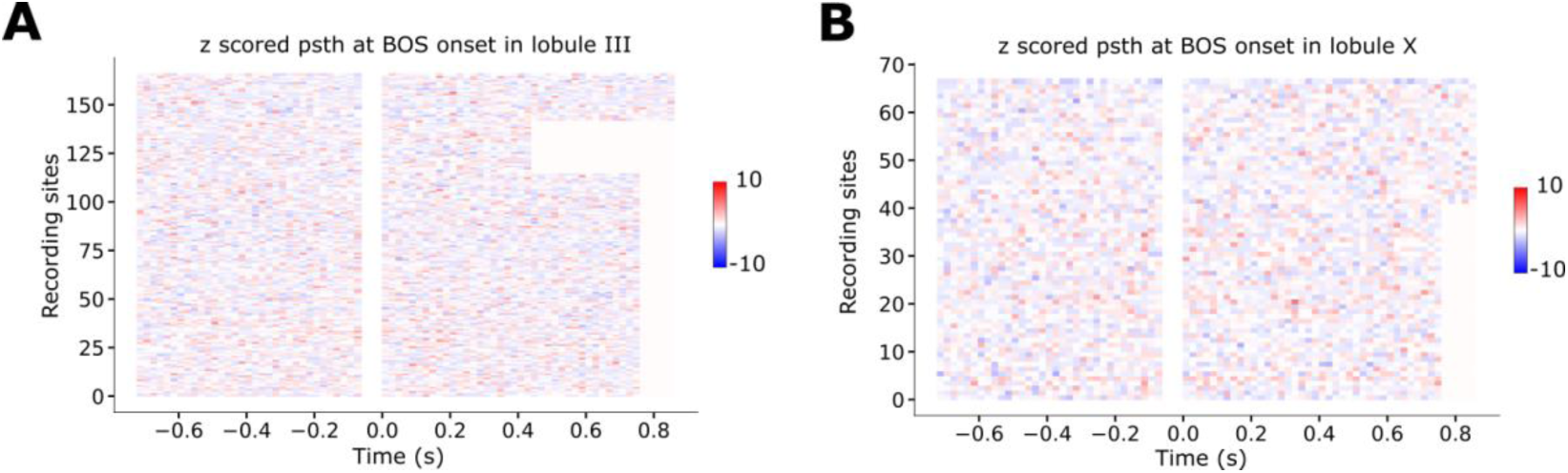
Response of the neuronal activity in cerebellar lobules to the onset of BOS. A: each line represents the colour coded z-scored psth of a recording site in the lobule III. Only the recording sites are selected for which z>2.5 or z<-2.5 after BOS onset. The green vertical dashed line represents the white noise onset. The bin for the psth is 20 ms. The colour code bar is on the right. Left: Baseline activity average over the same number of trials as BOS playbacks. Right: Average activity during BOS playback. B: same as A but for the lobule X.

**Supplementary figure 7:**
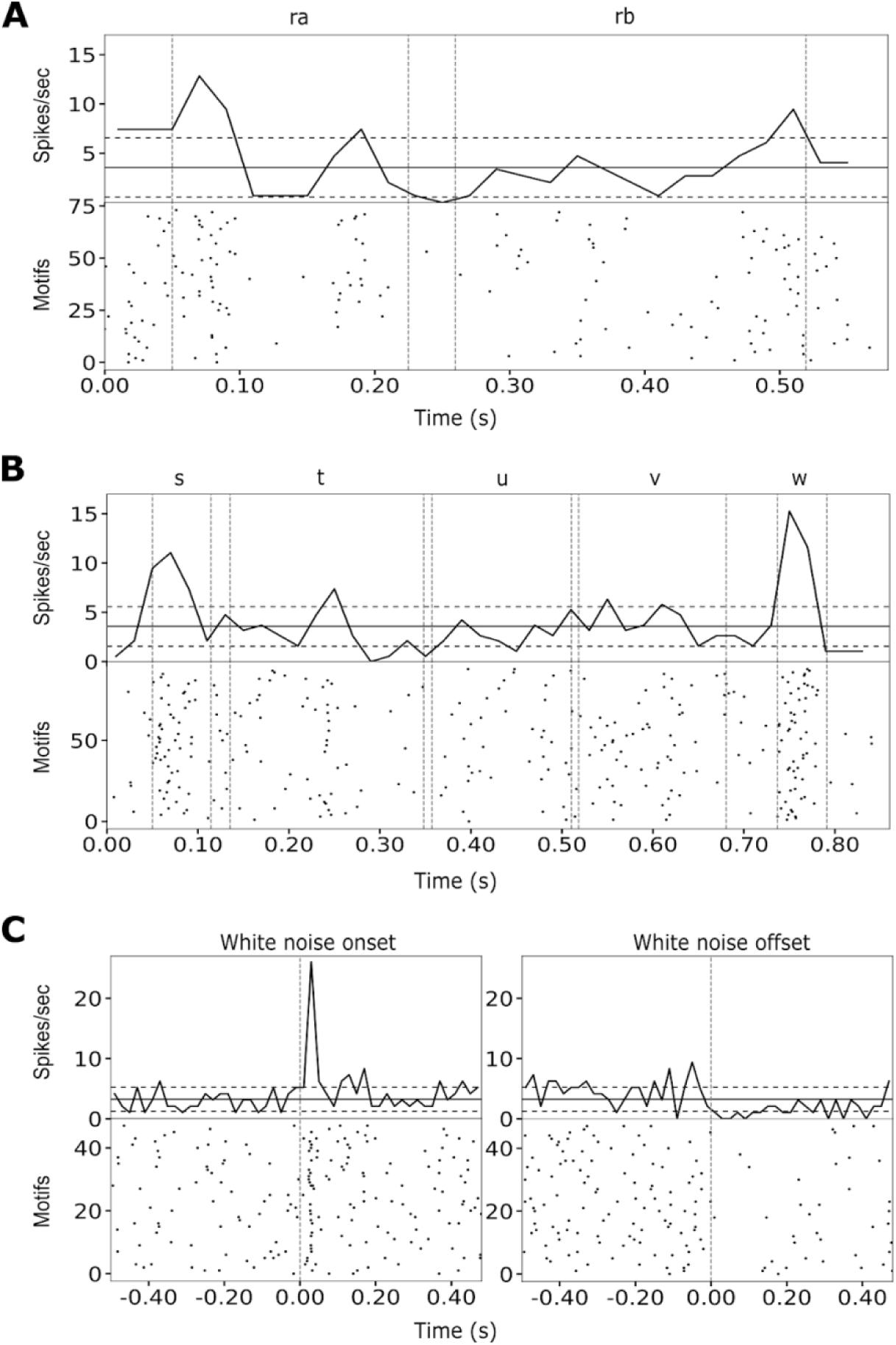
A: Same neuron and legend as figure 4,C but for reversed BOS: ra stands for reversed a, rb for reversed b. B: Same neuron and legend as figure 4,C but for other conspecific song. The conspecific song motif consists of the syllables s,t,u,v,w. C: Same neuron and legend as figure 4,C but for White Noise.

**Supplementary figure 8:**
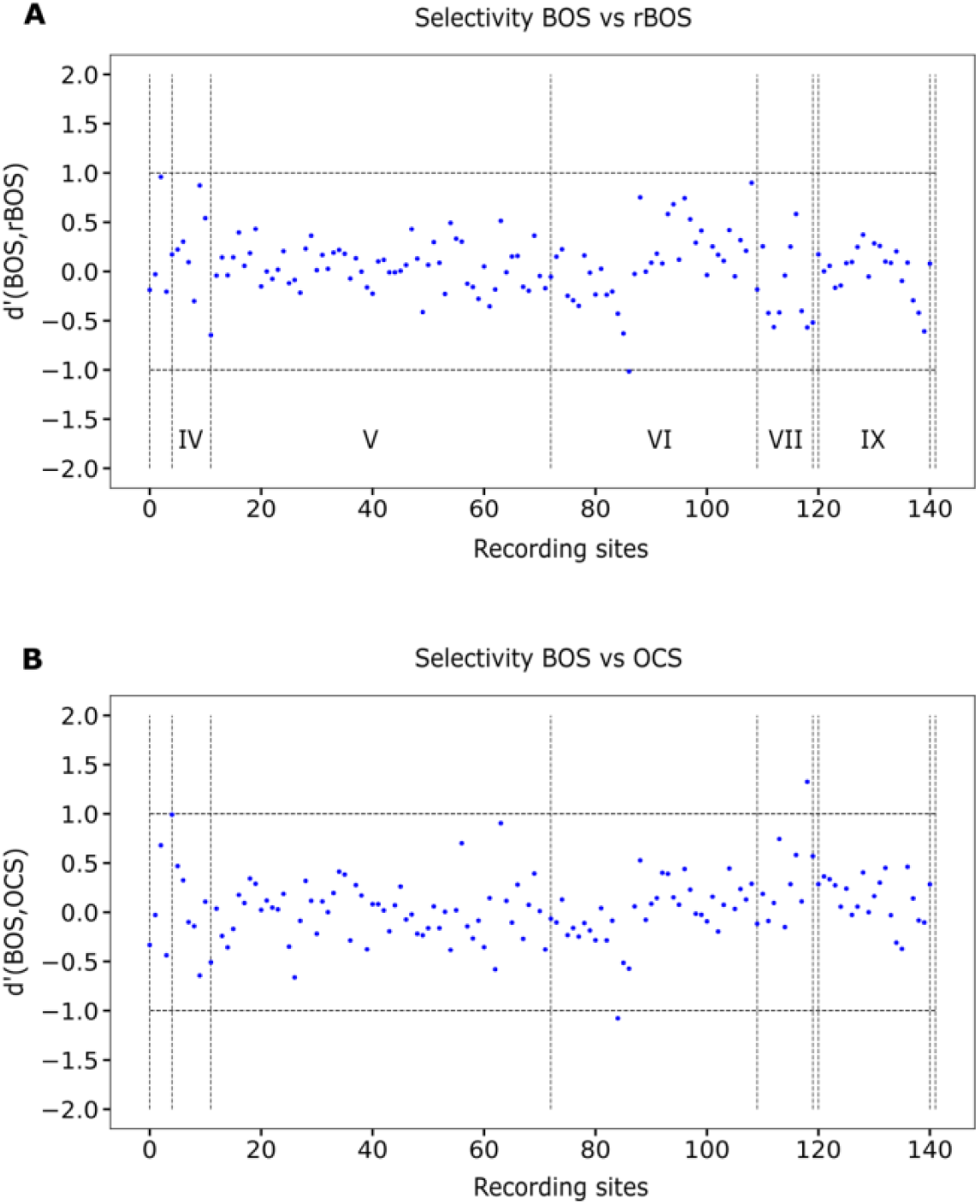
Selectivity of cerebellar neurons in awake recordings. A: Each dot represents the selectivity of the neuronal activity of the recording sites to BOS vs reverse BOS stimuli measured by d’. The horizontal dashed lines at d’=1/-1 represent the level of significance. d’>1 means the neuron is more selective to BOS than reverse BOS or conspecific song. d’<-1 means the neuron is more selective to conspecific song or reverse BOS than to BOS. The recording sites are sorted by lobule: outside cb, below the lobules, lobule IV, V, VIb, VII, VIII, IX, X. B: Each dot represents the selectivity of the neuronal activity to BOS vs reverse BOS stimuli measured by d’. The measure of selectivity d’ was proposed in (Solis & Doupe, 1997) and is a normalized difference of the mean firing rate during the two types of audio stimuli.

**Supplementary figure 9:**
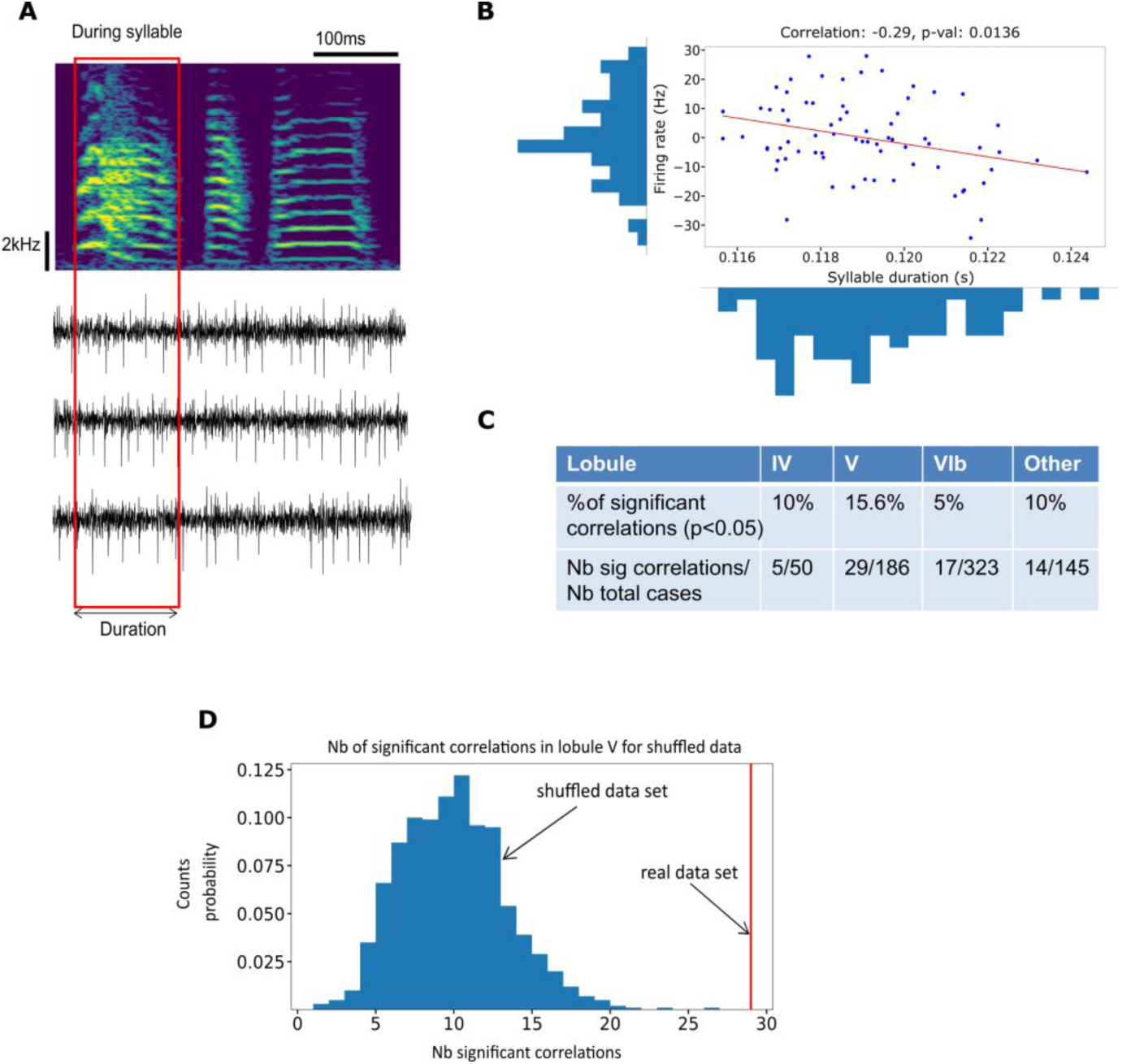
Correlations between the firing rate and syllable duration. A: Principle of measuring the correlation between the duration of a syllable and the firing rate during the syllable production. Example spectrogram (top) and neuronal firing (bottom) for 3 renditions of the motif. B: Example of a significant correlation between a multiunit neuronal activity and the syllable duration. C: Proportion of cases (syllable x recording site) featuring a significant correlation between the firing rate and the syllable duration in different lobules. D: Bootstrap analysis showing the significance of the proportion of significant correlations for the lobule V.

**Supplementary figure 10:**
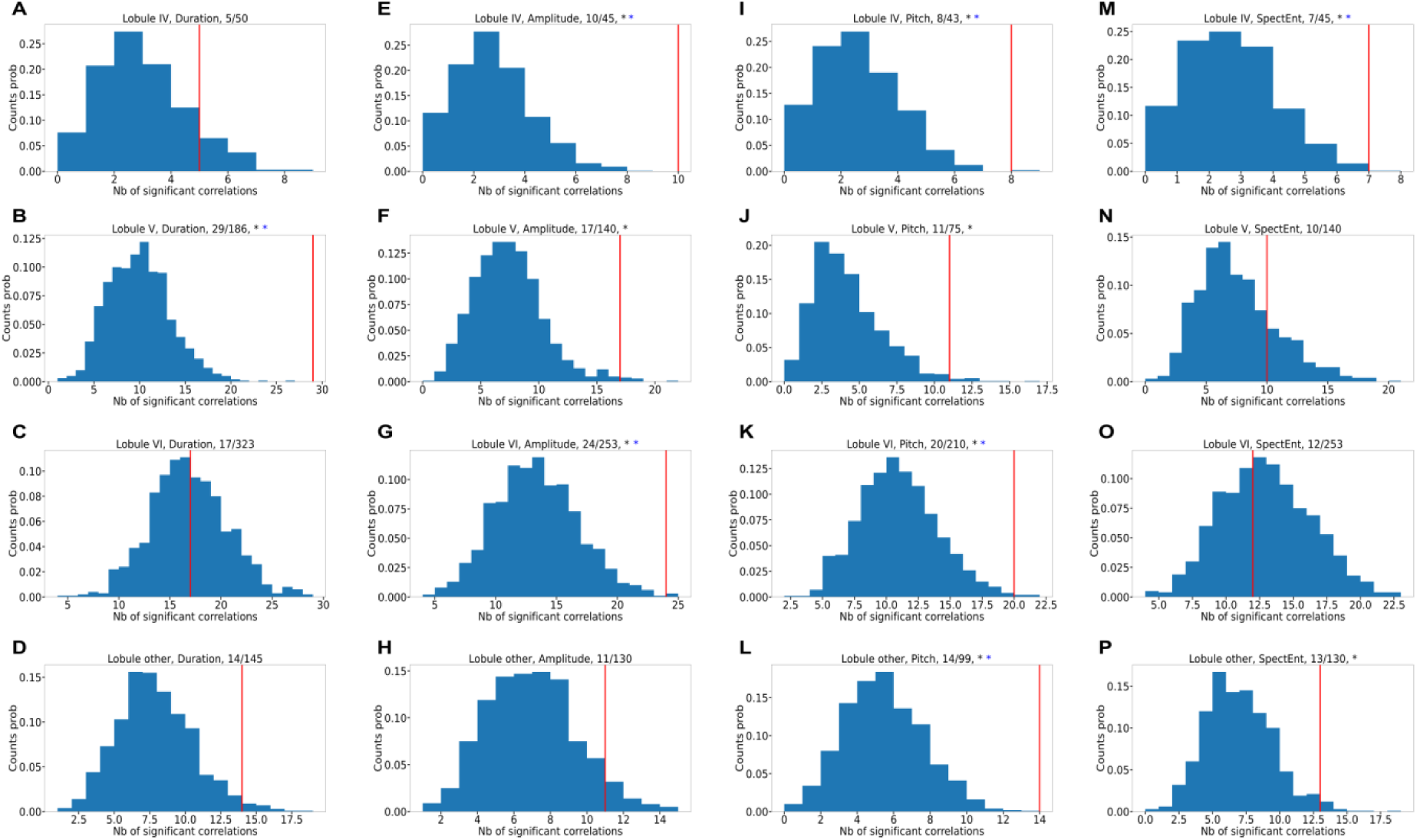
Bootstrap analysis for correlation between duration and detrended firing rate per lobule, pitch and firing rate in premotor window, amplitude and firing rate in premotor window and spectral entropy and firing rate in premotor window. The bootstrapped distribution of correlation coefficients is shown in blue. The number of significant correlations found in the real, non-shuffled data set is shown as a red vertical line. A, B, C, D: Bootstrap analysis for the correlation between the duration of a syllable and the activity during syllables for the lobules IV, V, VI and other lobules. E, F, G, H: Bootstrap analysis for the correlation between the amplitude of a syllable and the activity during a premotor window of the syllable for the lobules IV, V, VI and other lobules. I, J, K, L: Bootstrap analysis for the correlation between the pitch of a syllable and the activity during a premotor window of the syllable for the lobules IV, V, VI and other lobules lobules. M, N, O, P: Bootstrap analysis for the correlation between the spectral entropy of a syllable and the activity during a premotor window of the syllable for the lobules IV, V, VI and other. The title of each subplot specifies the lobule and feature for which the bootstrap analysis was conducted as well as the number of (syllable x recording site) cases with significant correlations among the total number of cases. The presence of an asterisk means that the proportion of (syllable x recording site) cases with significant correlations is significant according to the bootstrap analysis. Black asterisk represents significance without Bonferroni correction. Blue asterisk with Bonferroni correction

**Supplementary figure 11:**
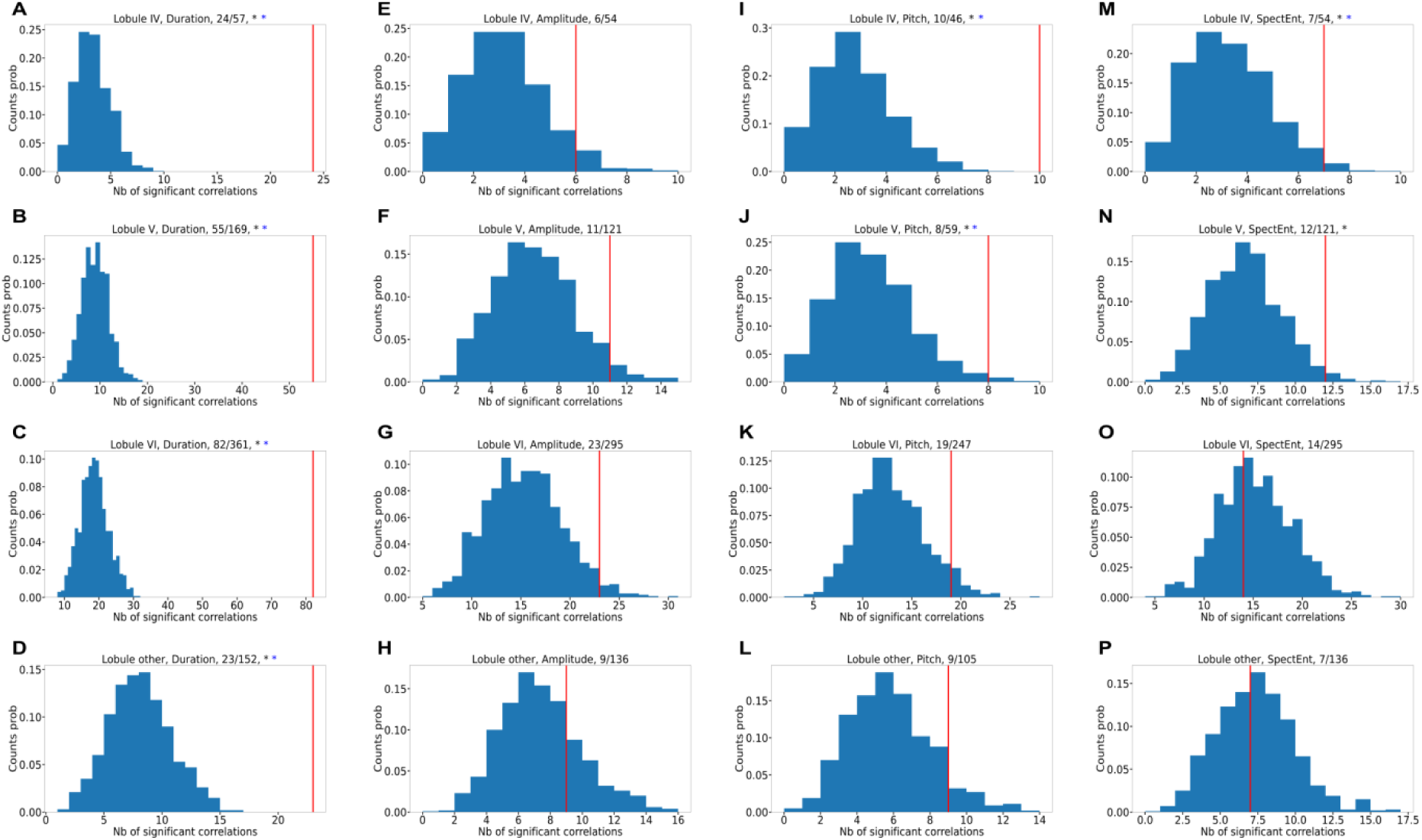
Same as supplementary figure 10 but without detrending the firing rate

**Supplementary figure 12:**
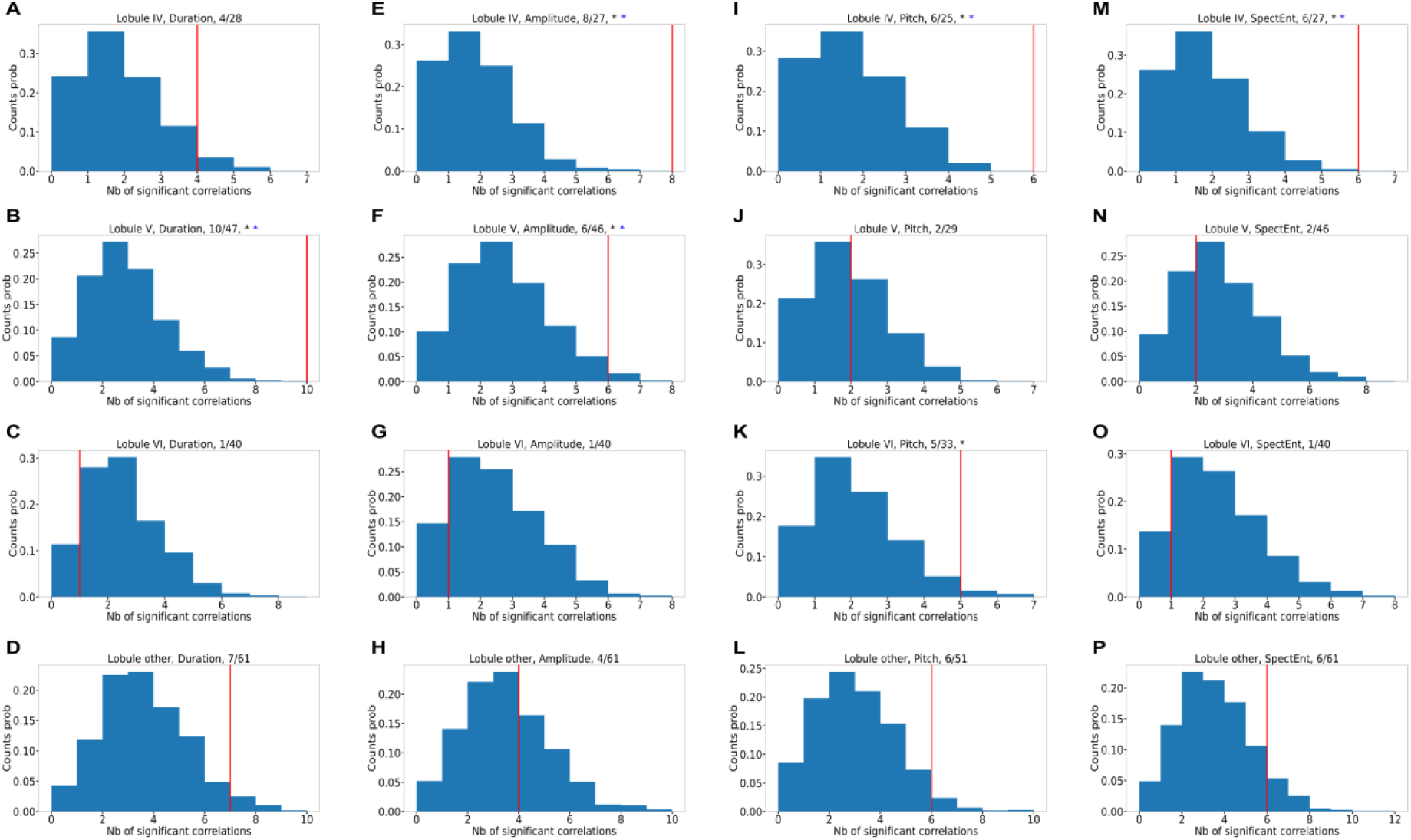
Same as supplementary figure 10 but for HF units.

**Supplementary figure 13:**
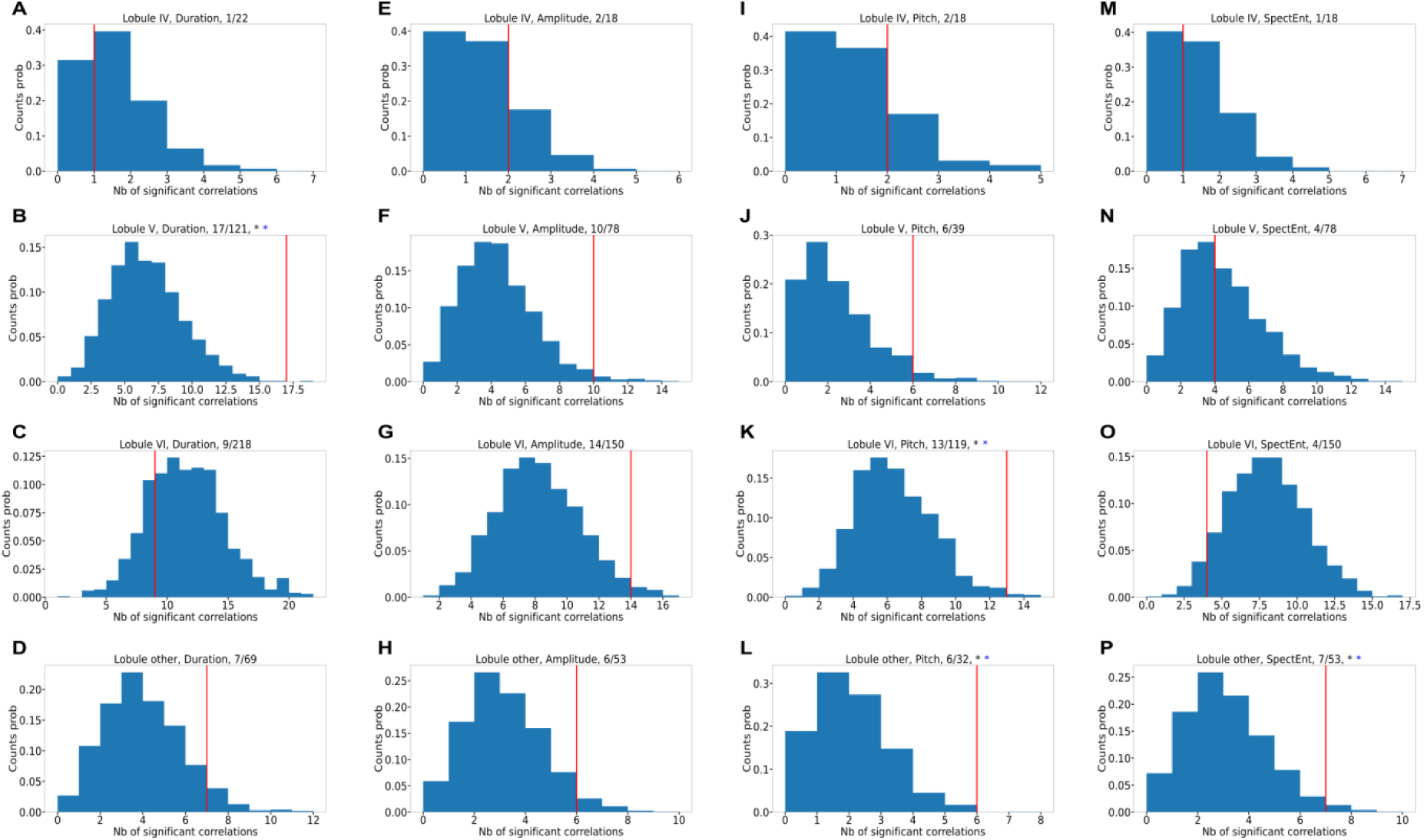
Same as supplementary figure 10 but for LF units. Correlations between neuronal activity and song features: We measured the correlation between the firing rate and the acoustic features (pitch, amplitude and spectral entropy, (Sober et al., 2008)) as well as the duration of the syllables (Suppl Fig 9-13). When measuring the correlation between a feature of a syllable and the firing rate, we compensated for the fluctuations or drifts of the firing rate on the long-time scales by detrending the firing rate (Suppl Fig 10, see Methods). We also measured the correlation between the duration of the syllables and the firing rate without detrending (Suppl Fig 11), since detrending in itself may induce spurious correlations (Harris, 2021). We only report correlations that were present whether detrending was applied or not to avoid artefactual detection of correlations. In the lobule V, the proportion of neurons that displayed significant correlation in their firing rate with the duration of the syllables amounts to 15% of all cases (syllablexrecording site, see example on Suppl Fig 9) and this proportion is unlikely to be found by chance (bootstrap analysis in Suppl Fig 9, 10 and 11). The significant correlations between syllable duration and firing rate in lobule V are present in the population of HF as well as LF cells (Suppl Fig 11B and Suppl Fig 12B). In the other lobules of the cerebellum, the proportion of significant correlations with the syllable duration is around 10% and is not significant when detrending the firing rate (Suppl Fig 9 and Suppl Fig 10). Among all other acoustic parameters, the pitch and spectral entropy of syllables was found to significantly correlate with the neuronal activities in lobules IV only (Suppl Fig 10 and 11), and when separating cell type we found significant correlation of these acoustic parameters with the HF firing rate only (Suppl Fig 12 and 13). Altogether, this correlation analysis should be interpreted carefully (Harris, 2021) but we believe it may point to a role of lobule V in tracking or modulating syllable duration fluctuations.

**Supplementary figure 14:**
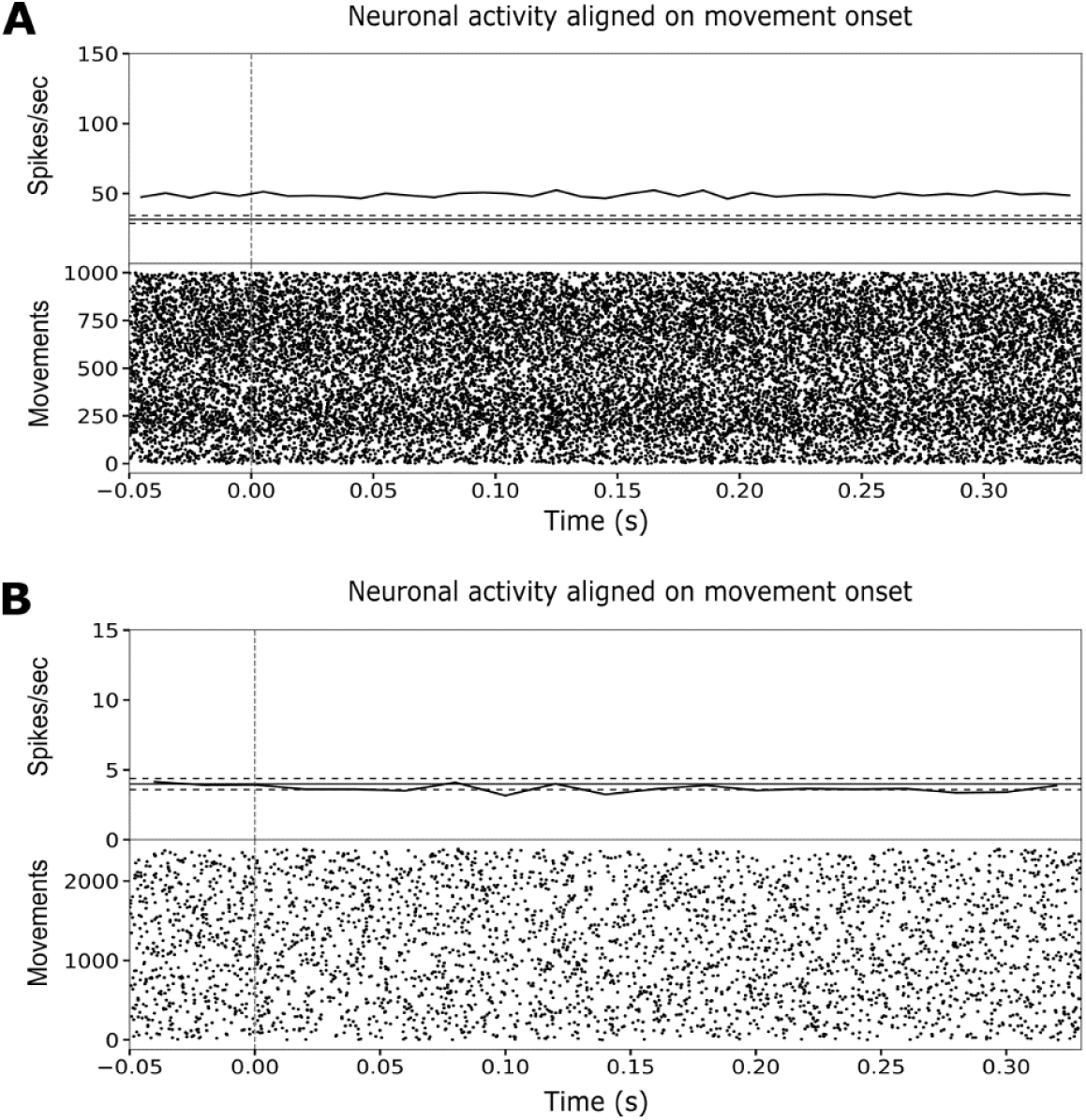
A: Activity of the neuron from figure 4 aligned on movement onset of the bird. Raster plot (bottom) and average firing rate PSTH (top, black). Each line of the raster plot corresponds to a movement of the body or head of the bird outside singing. Time 0 corresponds to the onset of the movement. B: Activity of neuron from figure 7 aligned on movement onset of the bird. Horizontal black lines represent the average firing rate and 95% confidence interval (dashed) during the same baseline period as in figures 4 and 7 respectively.

**Supplementary figure 15:**
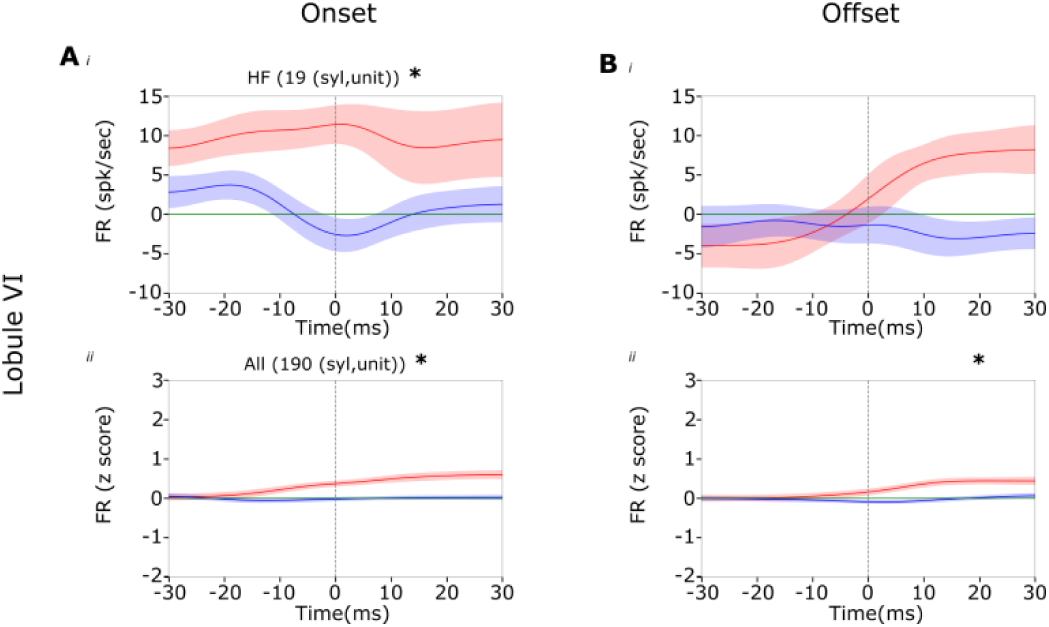
Same figure 7,F,G but for lobule VI.

**Supplementary Table 1:**
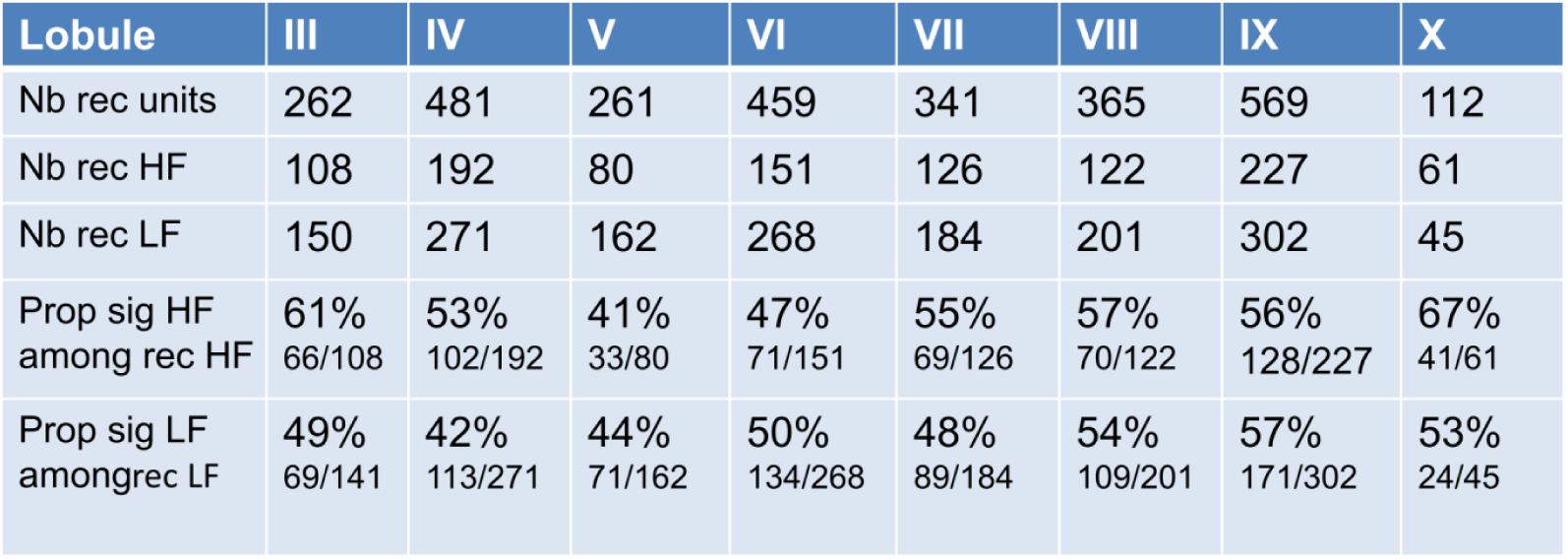
Total number of units recorded in anesthetized birds. Significant units are units with z scored firing rate during BOS above 2.5. HF are high firing single units. LH are low firing single units.

| Lobule | III | IV | V | VI | VII | VIII | IX | X |
| --- | --- | --- | --- | --- | --- | --- | --- | --- |
| Nb rec units | 262 | 481 | 261 | 459 | 341 | 365 | 569 | 112 |
| Nb rec HF | 108 | 192 | 80 | 151 | 126 | 122 | 227 | 61 |
| Nb rec LF | 150 | 271 | 162 | 268 | 184 | 201 | 302 | 45 |
| Prop sig HF among rec HF | 61%<br>66/108 | 53%<br>102/192 | 41%<br>33/80 | 47%<br>71/151 | 55%<br>69/126 | 57%<br>70/122 | 56%<br>128/227 | 67%<br>41/61 |
| Prop sig LF amongrec LF | 49%<br>69/141 | 42%<br>113/271 | 44%<br>71/162 | 50%<br>134/268 | 48%<br>89/184 | 54%<br>109/201 | 57%<br>171/302 | 53%<br>24/45 |

**Supplementary Table 2:** Alignment of motor and audio related activity in all recorded lobules of the cerebellum. Nb_recs is the number of recorded units during singing and BOS playback. Nb_sig is the number of recorded units with a significant alignment between singing and BOS evoked activity. HF and LF indicate the number of HF and LF units with significant alignments of singing and BOS evoked activity.

| Lobule | III | IV | V | VI | VII | VIII | IX | X |
| --- | --- | --- | --- | --- | --- | --- | --- | --- |
| Nb_recs | 0 | 12 | 71 | 72 | 10 | 3 | 19 | 2 |
| Nb_sig | 0 | 0 | 11 | 2 | 0 | 0 | 1 | 0 |
| HF | 0 | 0 | 2 | 1 | 0 | 0 | 0 | 0 |
| LF | 0 | 0 | 5 | 1 | 0 | 0 | 1 | 0 |

